# Engineered Living Materials based on Adhesin-mediated Trapping of Programmable Cells

**DOI:** 10.1101/734350

**Authors:** Shuaiqi Guo, Emilien Dubuc, Yahav Rave, Mick P.A. Verhagen, Simone A.E. Twisk, Tim van der Hek, Guido J.M. Oerlemans, Maxime C.M. van den Oetelaar, Laura S. van Hazendonk, Mariska Brüls, Bruno V. Eijkens, Pim L. Joostens, Sander R. Keij, Weizhou Xing, Martijn Nijs, Jetske Stalpers, Manoj Sharma, Marieke Gerth, Roy J.E.A. Boonen, Kees Verduin, Maarten Merkx, Ilja K. Voets, Tom F.A. de Greef

## Abstract

Engineered living materials have the potential for wide-ranging applications such as biosensing and treatment of diseases. Programmable cells provide the functional basis for living materials, however, their release into the environment raises numerous biosafety concerns. Current designs that limit the release of genetically engineered cells typically involve the fabrication of multi-layer hybrid materials with sub-micron porous matrices. Nevertheless the stringent physical barriers limit the diffusion of macromolecules and therefore the repertoire of molecules available for actuation in response to communication signals between cells and their environment. Here, we engineer a first-of-its-kind living material entitled ‘<u>P</u>latform for <u>A</u>dhesin-mediated <u>T</u>rapping of <u>C</u>ells in <u>H</u>ydrogels’ (PATCH). This technology is based on engineered *E. coli* that displays an adhesion protein derived from an Antarctic bacterium with high affinity for glucose. The adhesin stably anchors *E. coli* in dextran-based hydrogels with large pore diameters (10-100 μm) and reduces the leakage of bacteria into the environment by up to 100-fold. As an application of PATCH, we engineered *E. coli* to secrete lysostaphin via the Type 1 Secretion System and demonstrated that living materials containing this *E. coli* inhibit the growth of *S. aureus*, including the strain resistant to methicillin (MRSA). Our tunable platform allows stable integration of programmable cells in dextran-based hydrogels without compromising free diffusion of macromolecules and could have potential applications in biotechnology and biomedicine.

## Introduction

Synthetic biology aims to design programmable cells that combine sensing and molecular computing operations with on-demand production of proteins that have a broad spectrum of therapeutic applications [1–4]. Engineered living materials (ELMs) integrate genetically engineered cells with free standing materials and represent a new class of environmentally responsive living devices with designer physicochemical and material properties [5–7]. Ideally, ELMs provide mechanical robustness to engineered cells, prevent their leakage to the environment and allow cells to be viable for extended periods of time. The containment of genetically-modified microorganisms (GMMs) within various materials has become a grand challenge for future synthetic biology applications [8]. To date, strategies for containing GMMs inside a living device are based on the physical confinement by multi-layer materials [9–11]. Hybrid micro-patterned devices combining layers of elastomer and microporous hydrogel enabled the exchange of information with surrounding environment via diffusion of chemical inducers and their sensing by GMMs while displaying high mechanical resilience [10]. Nevertheless, the low porosity of the physical barriers mitigating bacterial escape also significantly hampers the diffusion of macromolecules, reducing the repertoire of synthetic biology applications to biosensing and release of small therapeutic molecules.

Programmable interactions between GMMs and their surroundings have recently gained interest as the design of multicellular structures such as engineered biofilms enables a higher control over shape and functionality of ELMs [12, 13]. Inducible secretion of fibrous proteins drives the emergence of molecular architectures supporting the artificial biofilms. Interestingly, such biofilms are self-regenerating and are able to perform advanced tasks such as multi-step enzymatic bioremediation, and are viable over weeks.

Inspired by this strategy, we propose to leverage the formation of programmable interactions between surface-displayed proteins on genetically engineered cells and hydrogel constituents to ensure cell containment within a living material while simultaneously allowing production and delivery of large therapeutic molecules. PATCH is based on engineered *E. coli* expressing a recombinant adhesion protein that stably anchors the cells to a hydrogel matrix, avoiding entrapment by stringent physical barriers (Fig.1a). Bacterial adhesins are modular cell-surface proteins that bind different ligands in a highly specific manner. While strategies for programmable cell-cell interactions via adhesins have been reported [14, 15], methods that allow adhesin-mediated cell-material interactions have not been described. We engineer *E. coli* bacteria to surface-display a Ca^2+^-dependent glucose-binding adhesin which has strong interactions with dextran-based hydrogels (Fig.1b and 1c for details). The engineered adhesin, referred to as *Mp*A, is derived from the *Marinomonas primoryensis* ice-binding protein (*Mp*IBP) which helps these bacteria to assemble into multi-species biofilms on ice [16–18]. Compared to conventional living material designs, our hydrogel-based material has large pores with diameters ranging from 10-100 μM, which substantially improves the diffusion of macromolecules through the matrix and bypasses complicated fabrication procedures. Our results reveal efficient containment of adhesin displaying bacteria with a retention factor of up to 100-fold higher compared to bacteria that lack the adhesin. As an application of PATCH, we anchor engineered *E. coli* that produce and secrete lysostaphin inside a dextran-based hydrogel (Fig.1d and e) and show successful inhibition of *S. aureus* growth including a Methicillin-Resistant (MRSA) strain. Taken together, our study demonstrates a programmable living material foundry with potential applications in biosensing and controlled delivery of therapeutic proteins.

**Figure 1:**
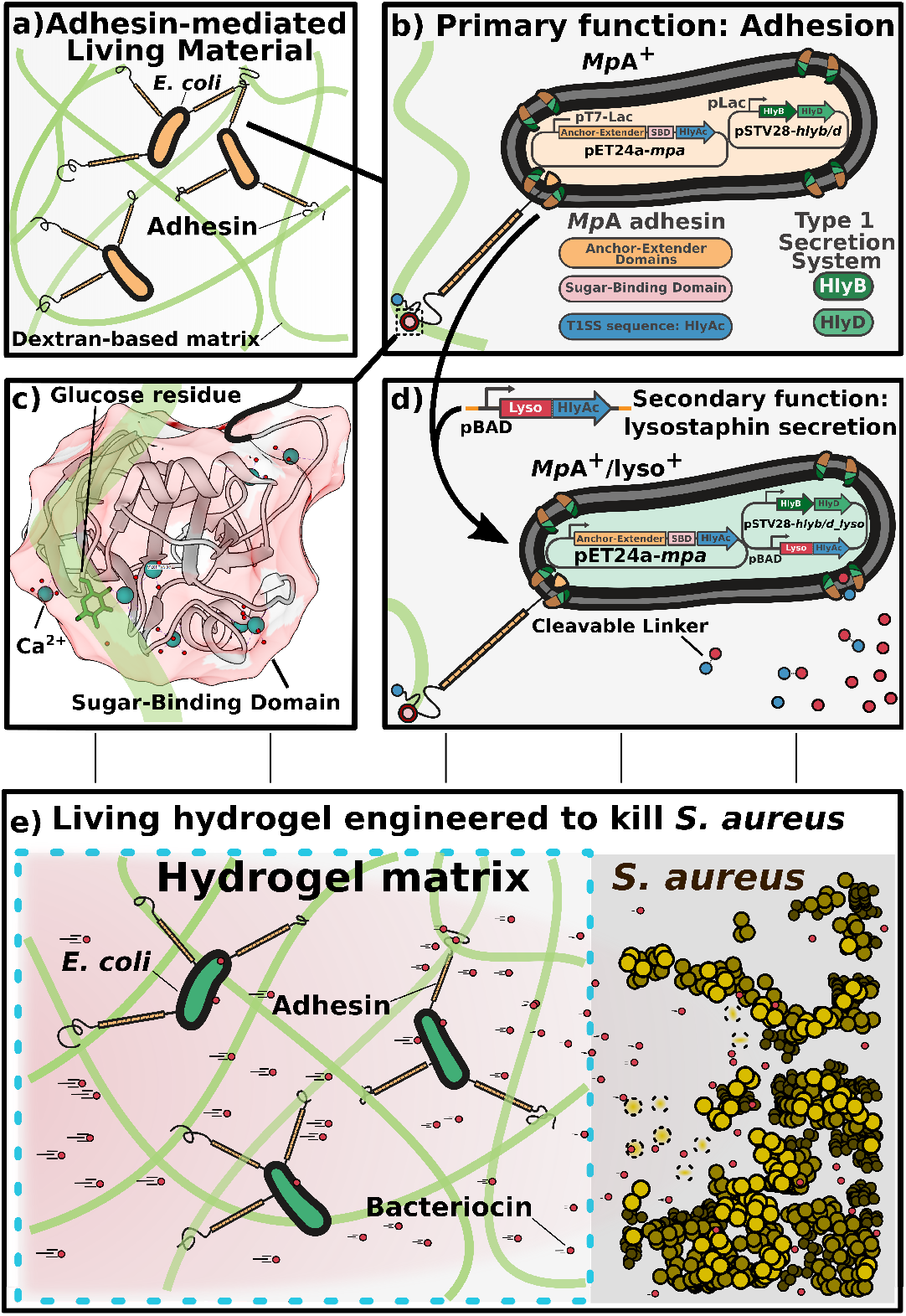
Overview of the design elements for PATCH. a) Escherichia coli (E. coli, orange ovals with black outlines) expressing a cell-surface adhesin (orange-black sticks) with a high affinity for glucose are retained inside dextran-based hydrogels. b) Cell surface display of the MpA adhesin is regulated by co-transforming E. coli with two expression vectors (MpA^+^; yellow). MpA was cloned into a pET24a vector (pET24a-mpa) under the control of a T7 promoter (pT7-Lac) inducible by IPTG. The engineered adhesin contains the membrane anchor (orange), extender (orange), and sugar-binding domains (SBD, pink circle) derived from MpIBP of the Antarctic bacterium, Marinomonas primoryensis. The 23-kDa C-terminal fragment of hemolysin protein A (HlyAc, dark blue circle) functions as a Type I Secretion Pathway (T1SS) sequence specific to E. coli, which was grafted to the end of MpA to promote its cell-surface-display. The other two E. coli T 1SS components HlyB (dark green) and HlyD (light green) were expressed from a pSTV28 vector (pSTV28-hlyb/d) under the control of a lac promoter (pLac) inducible by IPTG. The T1SS outer-membrane component (TolC) constitutive to E. coli BL21 cells is colored brown. c) X-ray crystal structure of the MpA sugar-binding domain (SBD), which is responsible for binding to the dextran component of the hydrogel matrix. Ca^2+^ are illustrated as large blue-green spheres, waters are indicated by small red spheres. The sugar-binding site is occupied by a glucose molecule (green backbone; stick representation). The Figure was rendered using UCSF Chimera [41]. d) E. coli (green; MpA^+^/lyso^+^) is re-engineered to surface-display the MpA adhesin and secrete the bacteriocin lysostaphin via the T1SS. HlyAc (small dark-blue circles) was added to the C-terminus of lysostaphin (small red circles) via a cleavable linker. The lysostaphin-HlyAc fusion construct was under the control of a pBAD promoter (inducible by arabinose) on the pSTV28 vector that also contains the pLac-HlyB/D. The modified vector is named pSTV28-hlyb/d_lyso. e) Schematics of the designed living material with anti-S. aureus activity. Left: Surface-expression of MpA allows E. coli to bind to the dextran matrix, thereby retaining the bacteria in the hydrogel (boundary marked by blue dashed lines). The engineered E. coli inside the hydrogel can secrete lysostaphin (small red circles), which diffuses freely to the exterior environment to inhibit the growth of S. aureus (gold spheres), including a methicillin-resistant strain (MRSA). Gold spheres with dashed boundaries indicate dead S. aureus cells killed by lysostaphin.

## Results and Discussion

### Synthesis and characterisation of dextran-based hydrogels

Dextran-based hydrogels are relatively simple to synthesize, robust in various shapes and highly biocompatible [19]. The glucose-based nature of the biopolymer makes dextran an ideal material for anchoring *E. coli* expressing *Mp*A. The dextran (500 kDa) polymer was functionalized by methacrylation to yield methacrylated dextran (Dex-MA) (Fig.S1), which was subsequently cross-linked into a covalent hydrogel via radical polymerization [20]. The resulting hydrogel is mechanically robust, and can be adapted into various shapes tailored for different applications. Further analysis of the wash solution of these hydrogels using NMR spectroscopy demonstrate the absence of toxic substances released from the gel, which makes it suitable for keeping bacteria viable and likely safe when applied onto human skin. Scanning electron microscope (SEM) analysis showed that the dextran-based hydrogel has an overall microporous structure with pore diameters ranging from 10-100 μm (Fig.S2). These large pores would allow unrestricted diffusion of macromolecules, as well as the propagation of native *E. coli* cells (2-5 μm), resulting in leakage of bacteria from the hydrogel in the absence of a surface-displayed adhesin (*vide infra*).

### Cell-surface display of MpA

To stably anchor *E. coli* to dextran-based hydrogel, we designed a new adhesion protein, *Mp*A, that binds glucose with high affinity and can be surface-displayed on *E. coli*. *Mp*A is derived from the giant adhesin *Mp*IBP (1.5 MDa) (Fig.2a) found on the cell surface of the Antarctic Gram-negative bacterium, *Marinomonas primoryensis*. *Mp*IBP binds its bacterium to ice and facilitates the formation of symbiotic biofilms with other microorganisms [17]. *Mp*IBP has a C-terminal signal sequence enabling its secretion by the Type I Secretion System (T1SS) specific to *M. primoryensis* (Fig.2a) [17]. At the N terminus, *Mp*IBP contains an “anchor module” that helps retain the protein to the cell surface by plugging the T1SS outer-membrane pore [17, 18, 21–23]. The protein further consists of 120 identical 104-aa tandem repeats that serve to project a set of ligand-binding modules away from the cell surface to interact with their target molecules, including various carbohydrates, proteins, and ice. The sugar-binding domain (SBD) of *Mp*IBP binds to glucose in a Ca^2+^-dependent manner [17, 18, 24], and is used in this study to allow engineered bacteria to bind to dextran-based hydrogels.

**Figure 2:**
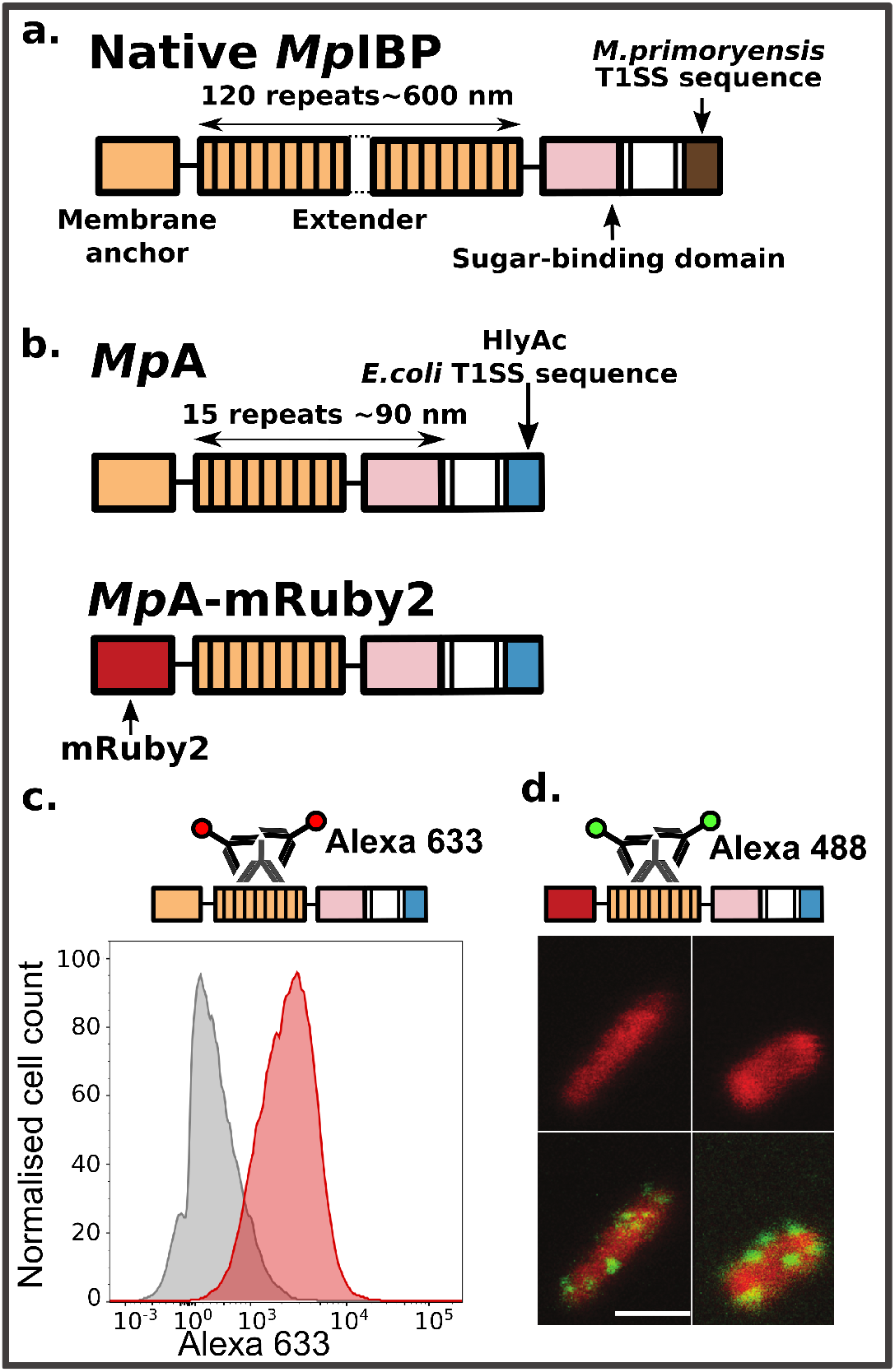
Construct design of the engineered MpA adhesins and its localisation to E. coli cell surface. a) Linear domain map of the native 1.5-MDa MpIBP with four of its functional regions illustrated as membrane anchor (orange), central extenders (orange repeats), sugar-binding domain (pink), and M. primoryensis T1SS sequence (brown). Dotted lines indicate the ~100 central repeats omitted in the figure. b) Linear domain maps of engineered constructs of MpA expressed and surface-displayed by E. coli. c) Immuno-detection of MpA by flow cytometry. Top schematic shows immunodetection of MpA through an antibody raised against its central extender domains. Histograms (bottom panel) illustrating fluorescence distribution of the immuno-detection experiments done with MpA^−^/lyso^+^ (grey) and MpA^+^/lyso^+^ (red) cells. d) Fluorescence localization after immunostaining (green) of MpA-mRuby2 (red) obtained using confocal microscopy. Top schematic shows the immunodetection of MpA through an antibody raised against its central extender domains. Bottom panels shows images obtained from confocal microscopy. The two MpA-mRuby2^+^ cells (red cells; top), and the detection of MpA on E. coli outer surface by immuno-staining with fluorescently-labelled secondary antibodies (Alexa 488, green patches on the cells; bottom). Scale bar indicates 2 μm, and is representative for all confocal images in this panel.

The extremely large molecular weight of *Mp*IBP prevents its use as a modular protein for cell-surface display in *E. coli*. We therefore introduced several modifications resulting in an engineered *Mp*IBP-based variant, *Mp*A, optimized for cell-surface display. First, we reduced the number of central *Mp*IBP repeats from 120 to 15, lowering the molecular weight by approximately 5-fold (to 330 kDa; Fig.2b). These 15 tandem extender domains can project the sugar-binding domain (SBD) approximately 90 nm away from the cell surface [17]. Next, the C-terminal domain (181 aa) of *Mp*IBP was replaced by a 23 kDa C-terminal sequence of the *E. coli* hemolysin protein A (HlyAc) (Fig.2b), which enables translocation of the protein to the cell surface via the Type I secretion pathway (T1SS) specific to *E. coli* [25–28]. The engineered *mpa* genes were placed in a pET24a vector. Since the genome of *E. coli* BLR (DE3) does not contain the T1SS machinery genes *hlyb* and *hlyd* required for secretion (pSTV28-*hlyb/d*; Fig.1b) [29], a pSTV28 vector carrying these two genes was co-transformed with the adhesin-pET24a vector (Fig.1b). We refer to bacterial cells co-transformed with these plasmids as *Mp*A^+^ cells and will use this notation throughout this article for other variants. Additional variants were created by replacing the N-terminal domains of *Mp*A with a fluorescent protein (*Mp*A-mRuby2^+^), which was used to validate the localization of the adhesin by confocal microscopy (Fig.2b). It has been reported that fluorescent proteins fold rapidly in the bacterial cytosol before being secreted by T1SS and their large size prevents full passage through the outer-membrane pore of the secretion system [27, 28]. Thus, mRuby2 serves as an “anchor” to hold *Mp*A to the cell surface, while the remaining parts of the protein protrude into the extracellular environments. As a control, we prepared *Mp*A^−^ cells by co-transformation with a pET24a plasmid that only encodes for HlyAc (pET24a-*hlyac*) and the pSTV28_*hlyb/d* vector. As detailed later in the paper, we also created *Mp*A^+^ cells capable of secreting the bacteriocin lysostaphin to target *Staphyloccocus aureus*. The resulting variant is referred to as *Mp*A^+^/lyso^+^.

Next, a combination of flow cytometry and confocal microscopy was used to validate the expression of *Mp*A on the outer surface of *E. coli*. *Mp*A^+^/lyso^+^ (cells capable of expressing both *Mp*A and lysostaphin proteins, Fig.1d, details below) and *Mp*A^−^/lyso^+^ control cells were incubated with rabbit anti-sera raised against the central repeats of *Mp*A. After unbound primary antibodies were removed by washing, cells were further incubated with a fluorescently-labelled (Alexa 488) secondary antibody (Fig.2c, top schematics). As expected, *Mp*A^+^/lyso^+^ cells (Fig.2c, red histogram) appeared approximately 5-fold brighter than that of the *Mp*A^−^/lyso^+^ cells (Fig.2c, grey histogram), validating the surface localization of the adhesin. Confocal microscopy studies using the *Mp*A-mRuby2^+^ variant revealed successful over-expression of the protein as evident by the evenly distributed red fluorescence inside of the cells. The surface display of this protein is demonstrated by the green immuno-fluorescence on the exterior of the cells. As the primary antibody targets the *Mp*A extender region consisting of 15 identical 104-aa repeats, it is likely that multiple antibodies bind to an individual adhesin protruding from the T1SS outer-membrane pore. This results in amplification of green fluorescence from the Alexa488 secondary antibodies, which appears as green clusters on the cell surface. These data are in agreement with the flow cytometry data (Fig.2d). In summary, we designed a novel *Mp*IBP-based protein for cell-surface display in *E. coli* and characterized the expression and localization of this protein using flow cytometry and confocal microscopy.

### MpA efficiently retains E. coli inside dextran-based hydrogels

To validate if surface expression of *Mp*A retains *E. coli* inside dextran-based hydrogels, we devised two experiments to quantify the leakage of bacteria through the hydrogel matrix into surrounding liquid or solid media. *Mp*A^+^ and *Mp*A^−^ cells were loaded into cubic-shaped hydrogels (Fig.S3), which were subsequently partially submerged in LB medium (Fig.3a). To estimate bacterial leakage from the gel to the surrounding medium, samples of the medium were harvested at different time points and plated for quantification by counting the colony forming units (CFUs). Samples taken immediately after loading of bacteria showed similar numbers (~ 10^2^) of *Mp*A^+^ and *Mp*A^−^ cells leaking into the medium. We hypothesize that this initial leakage originates from the large pore size of the hydrogel matrix resulting in a portion of loaded bacteria directly flowing through the gel without binding to dextran. After unbound cells were removed by refreshing the medium, very few *Mp*A^+^ bacteria were present in the medium outside of the hydrogel in the next 24 h (app. 30 CFUs). In sharp contrast, up to 100-fold *Mp*A^−^ cells were present in the medium (app. 2 × 10^3^ CFUs) (Fig.3b). SEM analysis was performed to gain visual insight into how bacteria interacted with the hydrogel. We observed that *Mp*A^+^ bacteria formed thin layers of microcolonies on the lamina of the hydrogel due to their strong interaction with the gel matrix (Fig.3c). In contrast, *Mp*A^−^ bacteria showed a significantly sparser distribution (Fig.S4), indicating a weaker interaction with the hydrogel, which was consistent with the results obtained from the leakage experiments.

**Figure 3:**
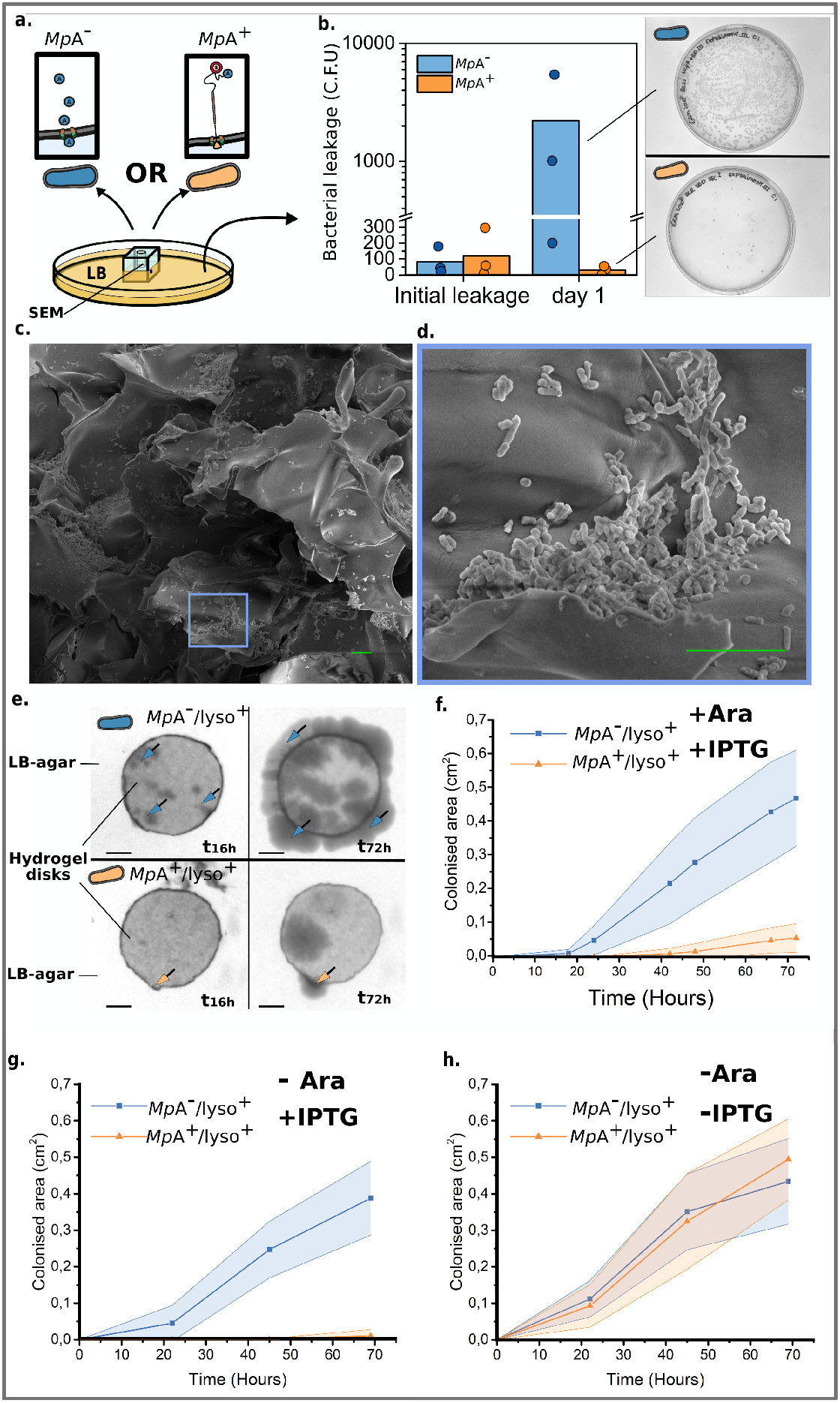
Characterisation of the MpA adhesin mediated retention of E. coli inside a dextran-based hydrogel matrix. a) Experimental design to quantify bacterial leakage from the hydrogel matrix into liquid medium. Leakage of MpA^+^ (orange) from a hydrogel cube was assessed by counting the colony forming units (CFUs) present in the LB medium surrounding the hydrogel, and compared with the leakage obtained from MpA^−^ cells (blue, negative control). b) Bacterial leakage was quantified immediately after bacterial loading (initial leakage) and over the following day. Right panel: representative images of plates at the same dilution are shown to illustrate the difference in leakage between MpA^+^ and MpA^−^ samples after 24 h (Day 1). Dots on the bar graph represent the results of individual experiments. c) Scanning electron microscopy images obtained from the dextran-based hydrogel inoculated with the MpA^+^ E. coli inside of the bacterial loading chamber (cross-section). d) Zoomed-in view of an area of (c), showing a bacterial microcolony of MpA^+^ formed on the lamina of the hydrogel matrix. Scale bars (green) in c) and d) indicate 10 μm. e) Representative images showing the migration of E. coli through the matrix of a disk-shaped hydrogel, which was assessed by estimating the surface area colonised by the bacteria (MpA^+^/lyso^+^ or MpA^−^/lyso^+^) on the outside of the hydrogel disk over time. Arrows indicate the presence of nascent colonies of E. coli (MpA^−^/lyso in blue) and (MpA^+^/lyso^+^ in orange). Scale bar represents 2 mm. f), g) and h) Quantification of the time-dependent migration of engineered E. coli (MpA^+^/lyso^+^ in blue and MpA^−^/lyso^+^ in orange) through the hydrogel matrix. Average area and standard deviation of the bacteria-colonized regions outside of the hydrogel were calculated from replicates of e) over 72 hours (n=4). f) Experiments performed in the presence of both inducing agents, IPTG and arabinose (Ara). g) Experiments performed in the presence of IPTG, but in the absence of Ara (n=3). h) Experiments performed in the absence of IPTG or Ara (n=3).

Next, we tested whether surface display of *Mp*A also inhibits migration of bacteria into a surrounding solid medium. Small disk-shaped dextran hydrogels (thickness: 0.75 mm; diameter: 8 mm) were fabricated. The hydrogel disk was then placed on a LB-agar plate, and *Mp*A^+^/lyso^+^ cells or *Mp*A^−^/lyso^+^ control (1 μL) were loaded to the center of the disk. Bacterial migration from the center of the hydrogel matrix was monitored for up to 72 hours (Fig.3e). We quantified the average colonised area outside the hydrogel disk based on four replicates (Fig.3f) and observed that after 72 hours *Mp*A^−^/lyso^+^ cells colonised a surface 8-fold larger compared to *Mp*A^+^/lyso^+^ cells (Fig.3d & 3e). Furthermore, uninduced *Mp*A^+^/lyso^+^ cells did not show significant differences in migratory behaviour compared to either induced or uninduced *Mp*A^−^/lyso^+^ control (-IPTG, Fig.3f-h). The expression and secretion of an additional protein, lysostaphin, which was induced by arabinose (details below, Fig.1d), did not appear to hamper *Mp*A-mediated retention of the engineered *E. coli* to the hydrogel matrix (Fig.3f). These results further consolidated the finding that *Mp*A surface display can substantially enhance the retention of bacteria inside dextran-based hydrogels through its sugar-binding capability.

### Application of PATCH - Living material engineered to kill MRSA

Having developed PATCH, we next assessed whether *Mp*A^+^ bacteria can be used to deliver functional proteins *in situ*. We focussed on engineering *E. coli* to secrete lysostaphin, a bacteriocin produced by *Staphylococcus simulans* to kill *S. aureus* [30]. The full-length lysostaphin is a 50-kDa preproenzyme with a modular domain architecture including a 36-aa N-terminal signal peptide followed by a region comprising 15 tandem repeats of 13-aa, and the peptidase and cell-wall-targeting (CWT) domains at the C terminus [31]. After its secretion from *S. simulans*, the signal peptide and the repetitive region of prepro-lysostaphin are sequentially cleaved to reveal its active form, with the CWT domain binding to its substrate on *S. aureus* cell surface and the peptidase domain responsible for the catalytic cleavage of the glycil-glycine bonds.

As the functional construct of lysostaphin was engineered for secretion via the T1SS by *E. coli*, the N-terminal *S. simulans* signal peptides and the tandem repeats of the preprolysostaphin were deemed redundant and therefore not included in the design (Fig.1b). Instead, the HlyAc sequence required by *E. coli* T1SS was fused to the C-terminus of lysostaphin via a linker consisting of a thrombin-cleavage site flanked by tetra-glycine motifs. Although the lysostaphin would in theory compete with the *Mp*A adhesin for the T1SS ducts, our data reveals sufficient secretion of functional lysostaphin (*vide infra*). The design of the linker motifs was intended such that the HlyAc domain can be selectively removed to release the catalytic form of lysostaphin. In addition, a 6X Histidine-tag was fused to the C-terminal end of the lysostaphin-HlyAc fusion protein for purification and detection purposes. The lysostaphin-HlyAc construct under the control of a pBAD promoter (inducible by arabinose) was incorporated into the pSTV28-*hlyb/d*, resulting in the pSTV28_*hlyb/d_lyso* vector (Fig.1d) [32, 33]. The pSTV28-*hlyb/d_lyso* was co-transformed with pET24a_*mpa* vector to create *Mp*A^+^/lyso^+^ bacteria. As a control, we also prepared cells lacking *Mp*A expression (*Mp*A^−^/lyso^+^), which have been transformed with the pET24a_*hlyac* and the pSTV28_*hlyb/d_lyso* vectors.

To reveal successful secretion of the 50-kDa lysostaphin-HlyAc, cell-free medium of *Mp*A^−^/lyso^+^ was analyzed by Western blotting with anti-histag primary antibodies. As expected, a band of approximately 50 kDa was detected for the intact lysostaphin-HlyAc (Fig.S5). In addition, a much more intense band was observed at approximately 25 kDa, which indicated the presence of HlyAc alone. This observation suggested that lysostaphin-HlyAc construct had been cleaved after secretion. Given the capability of lysostaphin to cleave penta-glycine and the presence of two tetra-glycine motifs linking lysostaphin and HlyAc, it is possible that the fusion protein underwent site-specific autolysis. Nevertheless, this would have no negative impact on the enzymatic capability of lysostaphin as it frees up its catalytic domains from HlyAc.

We quantified the efficacy of the secreted lysostaphin-HlyAc against *S. aureus*, using an adapted version of Kirby-Bauer (disk diffusion) test (Fig.4a) [34]. We applied cell-free medium of *Mp*A^−^/lyso^+^ bacteria to Mueller-Hinton agar plates pre-streaked with *S. aureus.* After an overnight incubation period, clear round exclusion zones formed where the medium was applied, indicating the effective inhibition of *S.aureus* growth (Fig.4b). In contrast, cell-free medium from the *Mp*A^−^ cells showed no activity against *S. aureus*, indicating that growth inhibition was solely caused by lysostaphin secretion. *Mp*A^−^/lyso^+^ cells were unable to inhibit the growth of a different Gram-positive pathogen, *Streptoccocus agalactiae*, demonstrating the specificity of lysostaphin. To further consolidate this result, we spotted living *Mp*A^+^/lyso^+^ cell cultures onto a Mueller-Hinton agar plate streaked with the notorious methicillin-resistant *Staphylococcus aureus* (MRSA), which is a major cause for hospital-acquired infections worldwide [35, 36]. Strikingly, we observed clear halo-like exclusion zones surrounding the *Mp*A^+^/lyso^+^ cells, indicating the inhibition of MRSA growth (Fig.4c). These results demonstrate that sufficient amount of functional lysostaphin was secreted even though the *Mp*A adhesin and lysostaphin compete for T1SS ducts. In sharp contrast, no such inhibition zones separated the *E. coli* cells without lysostaphin-producing capability (*Mp*A^−^ cells) and MRSA. Consistent with the results above, *Mp*A^+^/lyso^+^ cells could not kill *S. agalactiae*. These results demonstrated the high specificity and promising potential of *in situ* lysostaphin secretion for treating antibiotic-resistant *S. aureus*-related infections.

**Figure 4:**
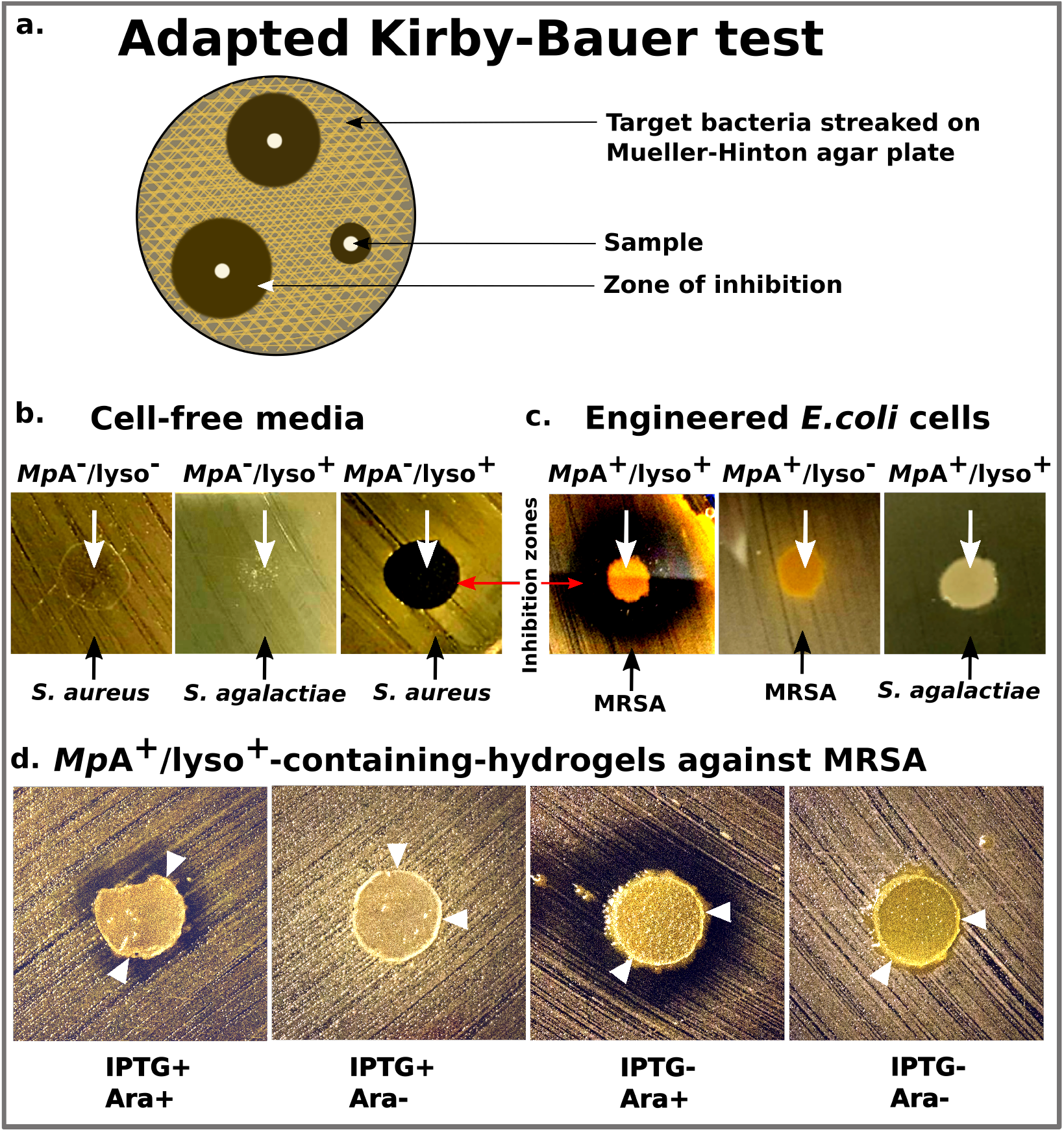
Anti-S. aureus activity of the novel E. coli-based living hydrogel. a) Schematic of the adapted Kirby-Bauer test for assessing the anti-S. aureus activity of lysostaphin-secreting E. coli. b) Evaluation of bactericidal activity of the cell-free media collected from MpA^−^/lyso^+^ or MpA^−^ cells against Staphylococcus aureus and Streptococcus agalactiae. White arrows indicate where the samples were applied. Black arrows indicate the target bacteria streaked on the plates. Growth Inhibition zones are marked by a red arrow. c) Evaluation of bactericidal activity of MpA^+^/lyso^+^ and MpA^+^ (no lysostaphin-producing capability) E. coli cells against Methicillin-resistant Staphylococcus aureus (MRSA) and S. agalactiae. d) Evaluation of bactericidal activity of the living hydrogels with engineered MpA^+^/lyso^+^ E. coli against MRSA. MpA^+^/lyso^+^ E. coli cells were spotted to the centres of the hydrogel disks, which were placed onto Mueller-Hinton agar plates streaked with MRSA. The images were captured after an incubation period of 96 hours. The adhesin- and lysostaphin-functionalities were controlled by the presence of IPTG and/or Arabinose (Ara), respectively. White arrowheads point to the edge of the hydrogel disks.

After demonstrating that *Mp*A^+^/lyso^+^ cells are able to secrete functional lysostaphin *in situ*, we next investigated whether hydrogels housing the engineered *E. coli* could kill MRSA, without leakage of GMMs. This would be a key step towards demonstrating the feasibility of this novel platform for different applications. As anticipated, hydrogel disks loaded with induced *Mp*A^+^/lyso^+^ *E. coli* showed no leakage into the surrounding Mueller-Hinton agar medium after 96 hours (Fig.4d). Remarkably, the bacteria-loaded hydrogel disks are also effective in killing MRSA, as exclusion zones developed around the periphery of the devices. This demonstrates free diffusion of functional lysostaphin through the hydrogel matrix to the exterior environments. In contrast, when *Mp*A expression was not induced (IPTG−), *Mp*A^+^/lyso^+^ cells were not restrained to the hydrogel, resulting in their migration into the exterior agar medium (Fig.4d, two panels on the right). When only lysostaphin expression was induced (IPTG−, Ara+), the inhibition zone was significantly larger than when both *Mp*A and lysostaphin were induced (IPTG+, Ara+). We reason that since the adhesin plugs a significant fraction of T1SS machineries, halting *Mp*A production resulted in higher secretion levels of lysostaphin. Furthermore, a control experiment in which lysostaphin expression was not induced (IPTG−, Ara−) resulted in the absence of an inhibition zone, which validates the responsiveness of our system towards arabinose induction (Fig.4d). Taken together, we have created a living hydrogel hosting adhesin-mediated self-retaining *E. coli*. As an application of this versatile platform, we demonstrated a prototype living material with anti-MRSA activity.

## Conclusions and outlook

One challenge that impedes the deployment of living functional materials for biomedical applications outside the lab is the requirement of sustaining viable cells while restricting their escape from the device. To the best of our knowledge, this study shows the first example towards confinement of GMMs by actively implementing adhesive interactions between microorganisms and a fully synthetic material. The basis of PATCH is in contrast with conventional methods for engineering living materials, which typically rely on the presence of a physical barrier provided by submicron matrices of multi-layer hybrid materials. Our work demonstrates one application of PATCH as a living material foundry for clearing pathogenic bacteria by *in situ* production and secretion of lysostaphin.

PATCH presents a versatile toolbox that can be tailored to the need of various applications such as biosensing and smart drug delivery in a deployable “Sense-kill” manner [37, 38], which opens the door for a wide range of future research. Identification of soluble *S.aureus* molecules capable of regulating the expression of lysostaphin would enhance the responsiveness and specificity of the current system. Implementing an alternative secretion system for lysostaphin might result in higher levels of *in-situ* drug delivery. Crucially, integration of the expression systems into the genomes of the engineered microorganisms will help diminish the spreading of antibiotic resistant genes to pathogens via the plasmids. To address further safety concerns, we envision that the platform could be adapted to other microorganisms that previously demonstrated their innocuity for humans, and were used in other animal models for synthetic biology applications, such as *E. coli* Nissle 1917 strain (EcN) [38] or *Bacillus subtilis* [39]. As such, this work constitutes an important methodological improvement in the field of living functional materials and living material foundries, and could have implications for therapeutic applications.

## Materials and Methods

### Gene synthesis of *Mp*A

A gene encoding a smaller version of *Mp*IBP (Genbank accession: ABL74378.1 and ABL74377.1) with only 15 central repeats was synthesized and purchased from GenScript (https://www.genscript.com). The gene was codon-optimized for expression in *E. coli*, and the central repeats were made as different as possible at the DNA level to reduce the chance of recombination. Next, the *M.primoryensis*-specific Type I Secretion System (T1SS) sequence near its C terminus (181 aa) was replaced by the *E. coli* T1SS sequence (last 213 aa of the Hemolysin A protein, HlyAc, also codon-optimized and synthesized by GenScript) to result in the *Mp*A adhesin (Fig.2b). A fluorescent variant of *Mp*A, *Mp*A-mRuby2, was created by replacing the membrane anchor (412 aa at the N terminus) of the adhesin with mRuby2 (Fig.2b). Each *Mp*A construct was placed between the *NdeI* and *XhoI* sites in a pET24a vector. Genes encoding for two other machinery proteins (*hlyb* and *hlyd*) of *E. coli* T1SS were placed between the *SacI* and *BamHI* sites in a pSTV28 plasmid as described [29]. The *hlyb/d* genes were PCR amplified from a pLG575 plasmid kindly provided by Professor Dr. Peter Sebo (Institute of Microbiology of the Czech Academy of Sciences).

### Flow cytometry

Approximately 10^6^ of either induced *Mp*A^+^/lyso^+^ or *Mp*A^−^/lyso^+^ cells were collected, blocked with 1 mg/mL bovine serum albumin (Sigma Aldrich) in a buffer containing 20 mM Tris-HCl pH 9, 150 mM NaCl and 5 mM CaCl_2_ (Buffer A), and incubated (1:100, 1h) with rabbit anti-serum raised against the central repeats of *Mp*A. After washing with buffer A (2X), cells were incubated with a goat anti-rabbit secondary antibody (Alexa 633, Invitrogen; 1:100, 1h) and then analysed by a Fluorescence Activated Cell Sorter (Aria III, BD Sciences). The stained cells were gated based on forward and side scattering. At least 10,000 gated events were analysed and plotted using the FlowJo software.

### Confocal microscopy

Cells were prepared in a similar way as those for flow cytometry experiments described above. *Mp*A-mRuby2-expressing cells were collected, blocked with 1 mg/mL bovine serum albumin (Sigma Aldrich) in Buffer A, and incubated (1:100, 1h) with rabbit anti-serum raised against the central repeats of *Mp*A. After washing with buffer A (2X), a secondary (goat anti-rabbit) antibody conjugated with a fluorophore (Alexa 488, invitrogen) was used to label the cell-surface-exposed *Mp*A-mRuby2. Samples were mounted on the stage of a confocal laser scanning microscope (Leica SP8) equipped with solid-state lasers (488 nm and 638 nm), a Plan APO 63× water immersion objective (NA 1.2) and hybrid detector. Fluorescent images were acquired using the Leica LAS X software.

### Synthesis of methacrylated dextran

Dextran T500 (500 kDa, Pharmacosmos) (4 g, 24.7 mmol glucopyranose unit) was dissolved in 36 mL dimethyl sulfoxide (DMSO) and degassed with argon for 15 min. After dissolving 800 mg 4-dimethylaminopyridine (DMAP, Sigma Aldrich), 2.4 mL glycidyl methacrylate (GMA, Sigma Aldrich) was added. The solution was stirred at room temperature for 44 h and the reaction was stopped by adding 6.6 mL of 1M HCl to neutralize the DMAP. The reaction mixture was transferred to SnakeSkin™ Dialysis bag (molecular weight cut-off: 10 kDa, Thermo scientific) and dialyzed against demineralised water at room temperature with changing water 3-4 times a day for 72 h. Next, the methacrylated dextran solution was lyophilized and stored at - 30 °C before use. The degree of substitution, which is the percentage of methacrylated glucose monomers, was calculated with the formula (x/1.04y)*100, where x stands for the average integral of the protons of the double bond, indicated with 22’ and 22“ in the spectrum, y stands for the integral of the anomeric proton, indicated with 4 in the spectrum, and 1.04 is the correction factor for the average 4% of α-1, 3 linkages in dextran. (see Fig.S1 for NMR spectra).

### Dextran-based hydrogel preparation

An aqueous solution of methacrylated dextran (5% w/v, 200 mg methacrylated dextran in 4 mL H_2_O) and ammonium persulfate (4.5 mg) was degassed with argon, stirred, and cooled in an ice bath for 15 min. From here on, all preparation was done at 4 °C in a cold room. After the addition of 5 μL tetramethylethylenediamine (TEMED), the solution was mixed vigorously for 4-5 seconds and poured into the mold for hydrogel formation via freezing at −20 °C.

### Mold fabrication for the cubic hydrogel

The device mold was fabricated in two steps. Firstly, a Poly(methyl methacrylate) (PMMA) mold was made using a CO_2_ laser (see Fig.S3a) and glued to a flat PMMA using an acrylic glue. Secondly, PDMS pre-polymer mixture was prepared in a 10:1 ratio (PDMS: curing agent) and poured over the mold. This was thermally cured in an over at 65 °C for 2h and carefully peeled off from the mold (see Fig.S3b). A side hole (Ø= 2 mm) was made at the center using metallic punches. In the side holes, glass needles were inserted before pouring in the hydrogel.

### Preparation of the cubic hydrogel for assessing the bacterial leakage

Activated methacrylated dextran solution was poured into a PDMS mold with a pin placed through the central hole from one side to the center of the cubic mold. Removal of the pin after the solidification of the pre-gel solution resulted in an empty chamber within the hydrogel for loading bacteria. The casted methacrylated dextran was kept frozen at −20 °C overnight, and later thawed at room temperature, washed with 20 mL Milli-Q, and allowed to swell for 5 min. Next, the gel was washed with 20 mL 0.05 M NaOH followed by 20 mL Milli-Q and stored at 4°C in a falcon tube with fresh Milli-Q.

### Hydrogel disk preparation

Activated methacrylated dextran was casted in a mold used for fabrication of acrylamide-based gels for SDS-PAGE (Mini PAGE mold, Bio-Rad). This resulted in the fabrication of a 0.75 mm-thick homogeneous sheet. After washing, the hydrogel disks were formed by punching off the sheet with a 8-mm hole puncher, and stored in fresh Milli-Q at room temperature.

### Cloning

Cloning procedures were performed following Gibson assembly method [32, 33]. Reagents and competent DH5a cells used for cloning were from the NEBuilder HiFi DNA Cloning Kit (Product E5520S New England Biolabs). Q5^®^ High-Fidelity 2X Master Mix was used for long PCRs (Product M0492SNew England Biolabs 240 County Road Ipswich, MA 01938-2732). The correct gene products were confirmed by DNA sequencing (BaseClear BV. Sylviusweg 74 2333 BE Leiden, The Netherlands, Tables S1 and S2).

### Bacterial culturing

Unless otherwise noted, all bacteria were grown at 37 °C. Cultures were grown overnight in LB supplemented with selective antibiotics from single colonies of freshly transformed cells. Overnight cultures were further diluted 100-fold in fresh LB supplemented with selective antibiotics. After 3 h incubation, cells were chemically induced (0 or 5mM arabinose, 0 or 0.5 mM IPTG) (OD_600nm_ = 0.8) and further grown for 3 h. Note timing is important for ensuring optimum protein expression levels and therefore functionality of the system.

### Bacterial leakage experiment with the cubic hydrogels (liquid LB medium)

Cubic hydrogels were soaked in LB medium supplemented with 30 μg/mL chloramphenicol, 100 μg/mL kanamycine, 5 mM CaCl_2_ and 0.5 mM IPTG for one hour and transferred to an empty sterile Petri-dish by tweezers with the opening of the bacteria-loading chamber facing upwards. Excess LB was pipetted out of the central chamber. Engineered *E. coli* (7 μL, OD_600_ nm is approximately 1) were pipetted into the central chamber of the cubic hydrogel placed in a sterile petri-dish. Next, 25 mL of medium was added to the plate gradually to roughly 80% (8 mm) of the height of the cubic hydrogel incubated at room temperature. The medium was collected at two different time points after the bacterial loading (10 min and 24 h) and plated (100 μl) for the quantification of bacterial leakage by CFU counting. The LB medium and Petri-dish were refreshed after each sample collection to prevent the over-growth of bacteria.

### Scanning Electron Microscopy

Dextran gels seeded with cells were first fixated in 4% formaldehyde for 20 minutes, and further kept in Milli-Q water. Samples were further sliced using a scalpel to show the cross-section of the bacterial loading chamber and sputter coated with a 9 nm-gold layer using an EMITECH K575X Peltier-cooled SOP Turbo sputter coater Dual. The gels were imaged at 5 kV in a FEI Quanta 3D field-emission SEM, equipped with an Everhart-Thornley secondary electron detector

### Bacterial leakage experiment with the hydrogel disks (LB-agar medium)

Prior to use, dextran-based hydrogel disks were soaked in LB medium supplemented with 30 μg/mL chloramphenicol, 100 μg/mL kanamycine and 5 mM CaCl_2_. Next, 100 μL of a solution containing chemical inducers (0 or 5mM arabinose, 0 or 0.5 mM IPTG) were spread on LB-agar plates supplemented with antibiotics and CaCl_2_. The surface of the LB-agar substrate was allowed to dry by incubating the plates at 37 °C for 30 min. Hydrogel disks were set on the agar-LB surface and quickly dabbed dry using clean, dust-free, paper tissue, and further incubated for 10 min at 37 °C. A drop (1 μL) of engineered *E. coli* culture was placed in the center of hydrogel surface on the LB agar medium within petri dishes. The migration of the engineered *E. coli* was tracked by imaging for 4 days. Pictures were analysed using ImageJ/Fiji [40]

### Antibacterial activity tests (modified Kirby-Bauer test)

100 μL master mix solutions containing various chemical inducers (0 or 5mM arabinose, 0 or 0.5 mM IPTG, 5 mM CaCl_2_) were spread onto 15 mL Mueller-Hinton agar plates, which were dried by incubating them at 37 °C. Further, a protocol of a standard Kirby-Bauer test was followed. Briefly, bacterial cultures of *Staphylococcus aureus* (methicillin resistant) or *Streptoccocus agalactiae* was diluted to OD_600nm_ of 0.4-0.6, and streaked across the Mueller-Hinton agar plates with a cotton swab before they were tested against various antibacterial treatments.

### Cell-free medium preparation

Approximately 3 hours after lysostaphin induction with 5 mM arabinose, 100 mL of engineered *E. coli* cell culture were spun down by centrifugation (4000 rpm) for 15 min at 4°C. The cell-free-medium was then harvested after the removal of residual cells by filtering (0.2 μm cut-off). To remove the antibiotics present in the growth medium, the cell-free-medium was buffer-exchanged into buffer A and concentrated approximately 30-fold using millipore spin cassettes (molecular weight cut-off: 10-kDa). The cell-free medium was further tested by applying a drop of 5 µxL onto the surface of the pre-streaked plates (see antibacterial activity tests).

### Living hydrogel with anti-MRSA activity

Dextran-based hydrogel disks were briefly dried using sterile cotton swab and then placed on the Mueller-Hinton agar plates streaked with methicillin-resistant *Staphylococcus aureus*. Prior to their infusion into the hydrogel disks, the engineered *E. coli* cells were washed to 2 times in LB to ensure complete removal of antibiotics (growth medium). Next, a drop (1 μL) of the induced *E. coli* cells was placed at the center of the dextran-based hydrogel disk surface. Plates were further incubated at 37 °C for up to 4 days and the effects of bacteriocin-producing living hydrogels were visualized by imaging with a DSLR camera (Canon 600D, 50mm lens). Pictures were analysed using ImageJ.

## Acknowledgements

The authors would like to acknowledge the financial support by STW-foundation, the EPSRC-NSFC Joint Research Project (No. 51461135005), the European Union (ERC-StG No. 635928 & 677313), the Dutch Science Foundation (NWO ECHO Grant No. 712.016.002 & NWO-VIDI grant No. 723.016.003), and the Dutch Ministry of Education, Culture and Science (Gravity Program 024.001.035) for funding this research. Dextran was kindly provided by Pharmacosmos A/S. We thank Ingeborg Schreur-Piet for her support for SEM imaging. We thank Prof. Peter Sebo for generously providing the plG575 plasmid for amplifying the HlyB/HlyD genes. We thank Bas Rosier for his help and valuable advises with the Figure 3b. We thank Prof. Luc Brunsveld for his important guidance and financial support while supervising the Tue 2018 iGEM team. Molecular graphics and analyses were performed with UCSF Chimera, developed by the Resource for Biocomputing, Visualization, and Informatics at the University of California, San Francisco, with support from NIH P41-GM103311.

