## Supplementary information for "Engineered Living Materials based on Adhesin-mediated Trapping of Programmable Cells"

**Table of contents.**

Fig.S1. ^1^H-NMR spectrum of methacrylated dextran.

Fig.S2. Scanning electron microscopy characterization of the microstructure of the dextran-based hydrogel matrix.

Fig. S3. Fabrication procedures of a cubic dextran-based hydrogel.

Fig.S4. Scanning electron microscopy images obtained from the dextran hydrogel inoculated with the *Mp*A^-^/lyso^-^ *E. coli* inside the bacterial loading chamber (cross-section).

Fig.S5. Expression and secretion of lysostaphin-HlyA demonstrated by SDS-PAGE and Western blotting.

Table S1. Primers and plasmid used in this study

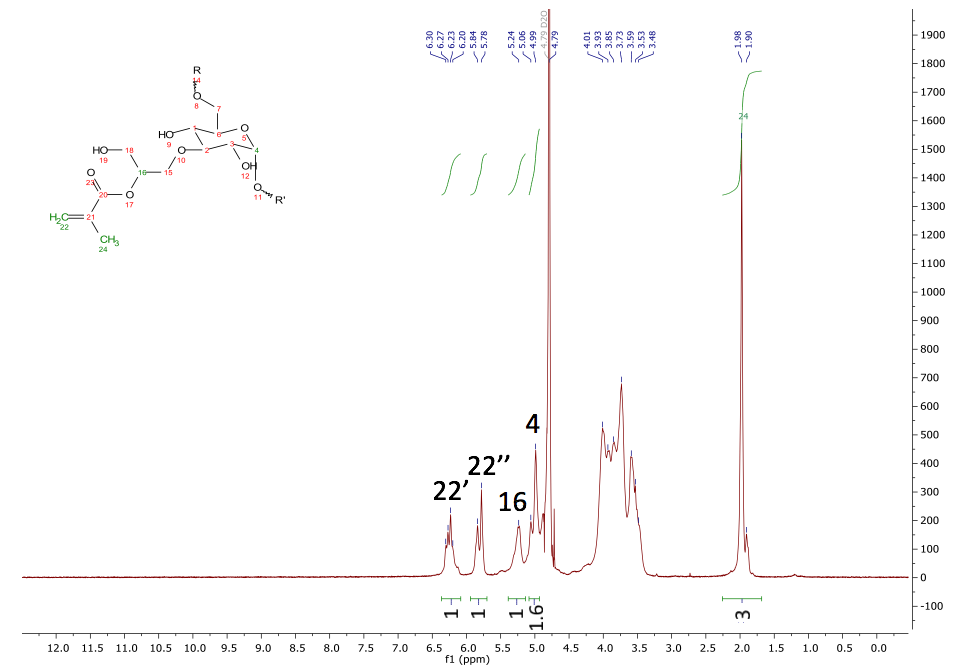

Fig.S1 ^1^H-NMR spectrum of methacrylated dextran with a degree of substitution of 60 prior to cross-linking. The degree of substitution, the percentage of glucose monomers that is methacrylated, was calculated with the formula (x/1.04y)*100, where x stands for the average integral of the protons of the double bond, indicated with 22’ and 22’’ in the spectrum, y stands for the integral of the anomeric proton, indicated with 4 in the spectrum, and 1.04 is the correction factor for the average 4% of α-1, 3 linkages in dextran. The assignment of the spectrum followed the method used by Plieva *et al.*^1^

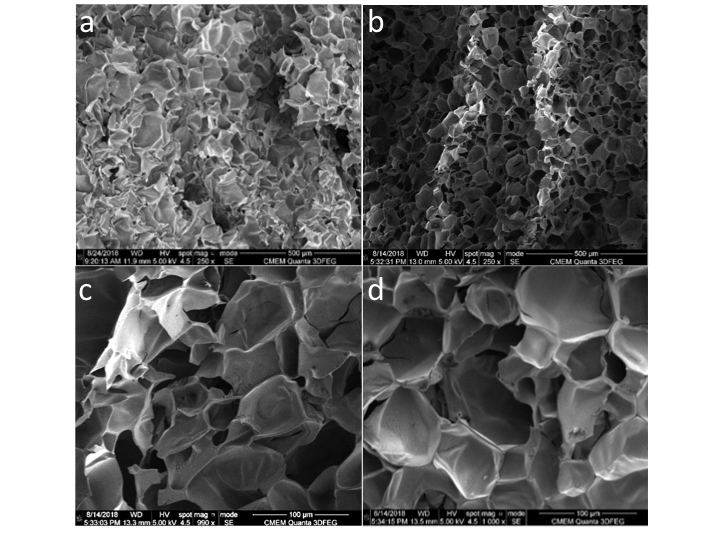

Fig.S2. Scanning electron microscopy characterization of the microstructure of the dextran-based hydrogel matrix. Overview of the microporous structure of the dextran-based hydrogel matrix (a and b). Zoomed-in views of the micro-porous structure of the dextran-based hydrogel matrix (c and d). The structure of the cross-linked dex-MA hydrogel is microporous with pore diameters ranging from 10 µm – 100 µm. Scale-bars are indicated.

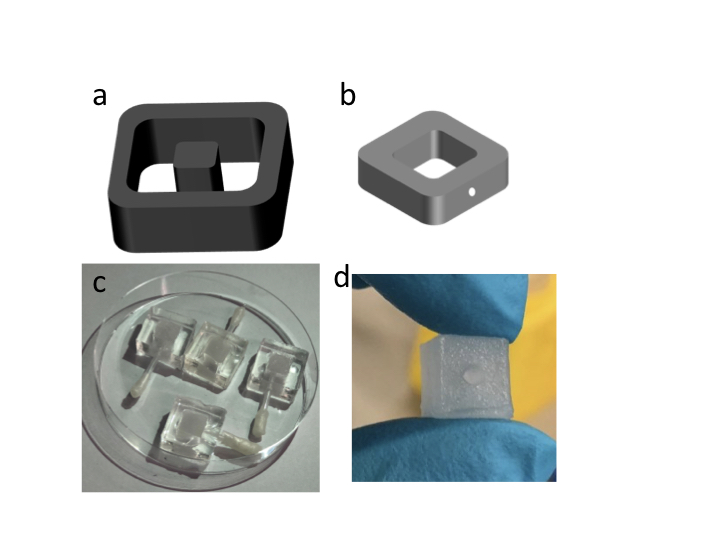

Fig. S3. Fabrication procedures of a cubic dextran-based hydrogel. a) Cartoon representation of poly (methyl methacrylate) (PMMA) mold fabricated using a CO_2_ laser with an outer wall dimension of 30 mm, an inner wall of 20 mm, and a central part of 10 mm. b) Polydimethylsiloxane (PDMS) mold peeled off from the PMMA mold, the white spot represents the hole with a diameter of 2 mm. A pin can be inserted half-way through the hole for making a bacteria-loading chamber at the center of the side walls as shown in c) and d). The inner dimension is 10 mm × 10 mm. c) Dextran-based hydrogel casted inside of the PMDS mold with a pin inserted half-way through the side-hole to create a central chamber for loading of bacteria. d) Close-up view of the cubic hydrogel with a central chamber for bacteria loading.

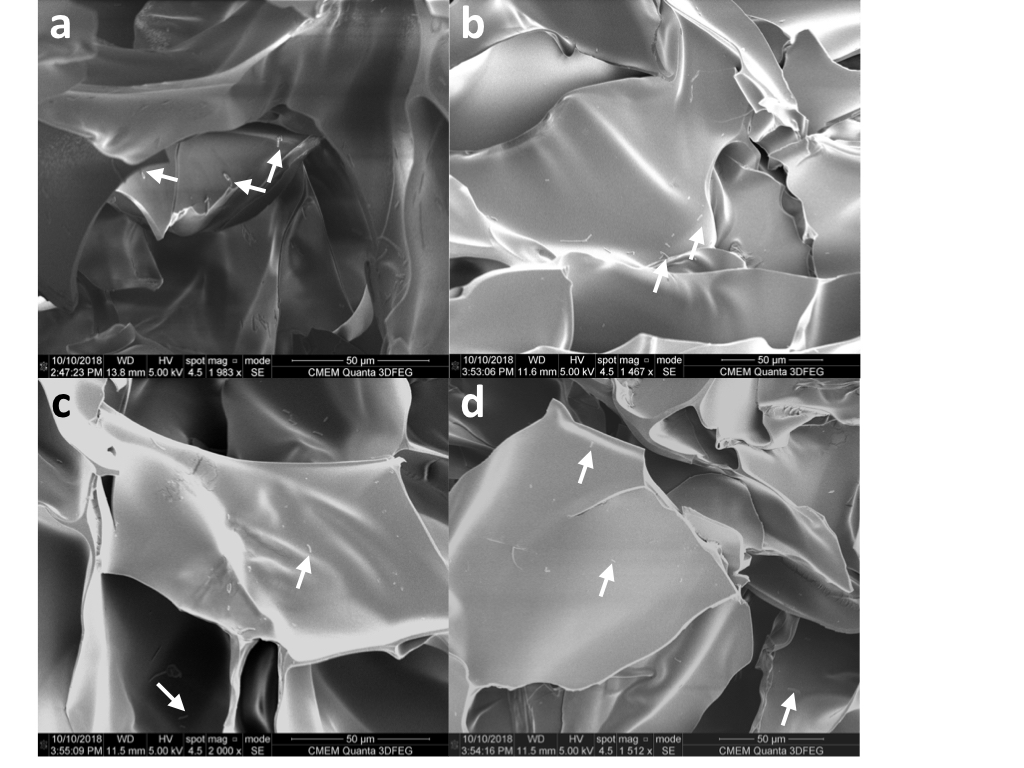

Fig.S4. Scanning electron microscopy images obtained from the dextran hydrogel inoculated with the *Mp*A^-^/lyso^-^ *E. coli* inside the bacterial loading chamber (cross-section). White arrows indicate the *Mp*A^-^/lyso^-^ *E. coli* on the hydrogel matrix. Scale-bars are indicated.

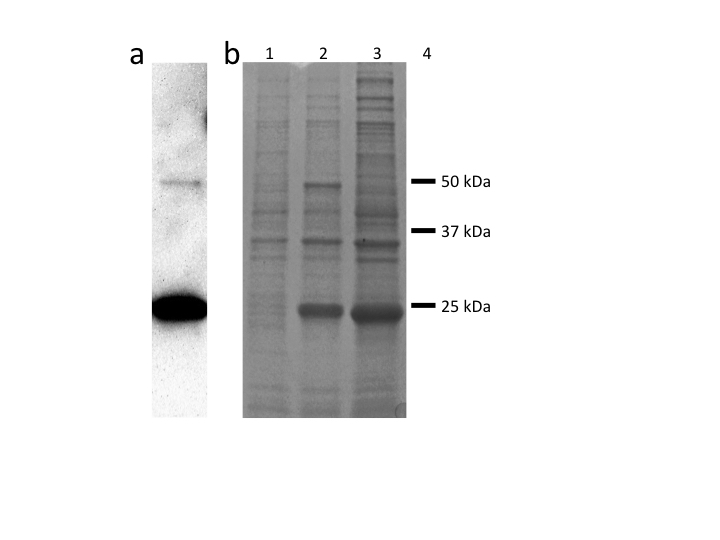

Fig.S5 Expression and secretion of lysostaphin-HlyA demonstrated by Western blotting and SDS-PAGE. a) The secretion of lysostaphin-HlyA and site-specific auto-proteolysis product are shown by Western blot. b) The expression of lysostaphin-HlyA (~ 50 kDa) is shown by SDS-PAGE. Lane 1: lysate of *Mp*A^-^/lyso^+^ uninduced for lysostaphin-HlyA expression. Lane 2: lysate of *Mp*A^-^/lyso^+^ obtained 3 h after induction for lysostaphin-HlyA expression. Lane 3: lysate of *Mp*A^-^/lyso^+^ induced for lysostaphin-HlyA expression obtained after an overnight growth period.

Table S1. Plasmids used in this study.

| **Plasmid** | **Promoter/Inducer** | **Source** |
| --- | --- | --- |
| pET24a | T7 Lac/IPTG | GenScript |
| pSTV28 | Lac/IPTG | TaKaRa |
| pBAD | pBAD/arabinose | Thermo Fisher Scientific |

Table S2. Primers and constructs used in this study.

| **Construct** | **Source** | **Identifier or Sequence** |
| --- | --- | --- |
| *Mp*A | Refs 16, 17, DNA synthesized by Genscript | See DNA sequence below |
| HlyAc | DNA synthesized by GenScript | See DNA sequence below |
| HlyB/D | PCR - amplified from a pLG575 plasmid kindly provided by Professor. Peter Sebo | See DNA sequence below |
| pSTV 28-hlyBD | Ref 29 | See Ref 29 |
| Lysostaphin-HlyA-His6 | This study | See DNA sequence below |
| pBAD-Lysostaphin (enzyme only)-HlyA-HisX6 REV (1) | This Study | 5’- ACAAGGGCCTTTGAGGTCATGGTATATCTCCTTCTTAAAGTTAAAAAACGG -3’ |
| pBAD-Lysostaphin (enzyme only)-HlyA-HisX6 FWD (2) | This study | 5’- CTTTAAGAAGGAGATATACCATGACCTCAAAGGCCCTTGT -3’ |
| pBAD-Lysostaphin (enzyme only)-HlyA-HisX6 REV (2) | This study | 5’- CAGTGGTGGTGGTGGTGGTGtgctgatgctgtcaaagtta -3’ |
| pBAD-Lysostaphin (enzyme only)-HlyA-HisX6 FWD (1) | This study | 5’- taactttgacagcatcagcaCACCACCACCACCACCACTG -3’ |
| pBADmod1-linker2-gamS REV | This study | 5’- ttagcaagagaatttcccatGGTATATCTCCTTCTTAAAGTTAAAAAACGG -3’ |
| pBAD FWD | This study | 5’- CTTTAAGAAGGAGATATACCatgggaaattctcttgctaaaa -3’ |
| pBAD REV | This study | 5’- CAGTGGTGGTGGTGGTGGTGtgctgatgctgtcaaagtta -3’ |
| pBADmod1-linker2-gamS FWD | This study | 5’- taactttgacagcatcagcaCACCACCACCACCACCACTG -3’ |
| pSTV 28-hlyBD FWD | This study | 5’-GTATCCGCTCATGAGACAATCACCATTATGTTCCGGATCTGCATCG -3’ |
| pSTV 28-hlyBD REV | This study | 5’- TCCCGGAGACGGTCACAGCTCAGGGCGCTTGTTTCGGCGT -3’ |
| pBAD-Lysostaphin-HlyA-His6 FWD | This study | 5’- ACGCCGAAACAAGCGCCCTGAGCTGTGACCGTCTCCGGGA -3’ |
| pBAD-Lysostaphin-HlyA-His6 REV | This study | 5’- AGATCCGGAACATAATGGTGATTGTCTCATGAGCGGATACATATTTGAA -3’ |

Nucleotide sequences of constructs used in this study. The nucleotide colors correspond to the color scheme in Fig. 1.

*Mp*A

ATGAGCGACAATCAGTTTCCGTTCGCCACCTTGGGCAATGCAATCGGCTTTATCACGAAGTTGGATGGTAGCGTCACCGTTCAGAGCATCGACGGCCAAGAACGCGTGTTGAAATTGGGCGACCCGATTTTCTTTGGCGAAACCGTGTTGACCGGTGGTTCCGGTTCTGTCACGATCGCGTTCGTTGATGGTACGGATGTCGTGATTGGTGGTGATTCCATCGTTGAAATGACCGATGAGATTTACAACACCGGCGATAACGAGGATCTGGTGGCAGACAGCTCTAGCGAGATTGATGCCCTGCAAAATGCAATCCTGGCAGGCGATGACCCGACGCTGATTCAAGACGCCCCGGCTGCCGGCAATACGCTGGCCGACCAGCAGCGCGTCGATGTTTCCATTGAACGTAACGATAATTCCGCCCAAGCTGGTTTCGGCGTGGACACCCAATCCAGCCTGCCGACCTACGGCTACGATACGGACAATGGTAATGGCGGCCAAGCGACCGAGCGCGAATACTCTGCTCCGAGCCTGAGCCGCACCTTGAACCAGTCCCCGCTGCTGATCAATCTGGATATCGATCCGGTTACCGGTGACTCTGTGATTAACGCAGCAGAAGCGGGTGGTACGGTGACCTTGACCGGTGTGGTGAACGGTGACGTTTTTAGCTCCGGTGTGGTTACGCTGGTTATCAATGGTGTGACGTACTCCACGAATGTGAATCCGAACGGCACGTGGTCTGTCTCTGTTGCAGGTTCTGATCTGAGCGCGGACTCCGATCGTATTGTTGATGCAAGCGTTGTGGTCACCAACGGTGCGGGTCAGCAAGGTACGGCTGATTCCACCGAGTCCTTTATCGTCAAGACGTCTAGCCGCGCTACCATCCGCGTTAATTCCATCACGTCCGACGACGTTGTTAATGCTGAGGAGAGCAATTCCACCATCACGGTTTCTGGCCGCGTCGGCTTGGACGCCTCCGCCGGTGATACGGTCTCTATGACGATCAATGGCACCCTGTACACCACCGTTGTGTTGGCTAACAAGACCTGGTCTGTGGGCGTGAGCGGTTCTGACTTGGCACAAGATAATTCTTTTCAGGTGTCCGTGACGGGTCAAGATTCTGCGGGTAATCCGTATGCAGGCACCACGACCAGCACCCATACGGTGGATACGTCTGCAGACGCGGGCACGGTGACCGTTAATGCTATTACCTCTGACGACGTGATCAACGCTAGCGAAGCGGCTGGTACCGTCGCCGTCTCTGGCACGGCGACGGGTGGCGATATTGCCGAAGGTGACACCGTTACGTTGGAGATCAACGGTGAGACGTACACCACGACCGTGGATGCGAATGGTGAATGGAGCGTGGATGTCGCGGGTAGCGATCTGGCTGCGGACACGGCTTTCGATGCTGTTGTTACGTCTTCCGATGCAGCCGGTAACACGGTTGATACCACCGGCTCTTCCACCCATACCGTCGACACGGAGGCAACCGCTGGTACGGTTACCGTGAATGCTATCACGTCCGATGATGTCATTAACGCATCTGAAGCAGCGGGCACCGTTGCCGTTTCCGGCACGGCAACCGGCGGCGATATCGCAGAGGGTGATACGGTGACCTTGGAAATTAATGGTGAGACCTACACCACGACGGTGGATGCCAACGGCGAATGGAGCGTTGATGTTGCAGGCAGCGACCTGGCCGCTGACACGGCGTTTGACGCTGTTGTGACCTCTTCTGATGCTGCAGGTAATACGGTTGATACGACGGGTTCCAGCACCCACACCGTTGATACCGAAGCTACGGCTGGTACCGTTACGGTTAACGCGATTACGAGCGATGACGTCATCAACGCGTCCGAAGCCGCTGGTACCGTTGCGGTTTCTGGTACCGCAACGGGCGGCGATATTGCCGAAGGCGACACGGTTACCTTGGAGATTAATGGCGAGACCTATACCACGACCGTCGATGCGAACGGCGAATGGTCTGTTGATGTGGCGGGTAGCGATCTGGCAGCTGACACGGCATTTGACGCCGTGGTTACGAGCAGCGATGCGGCCGGTAATACGGTGGACACGACGGGTTCCTCCACGCACACGGTTGATACGGAGGCGACGGCGGGCACCGTTACCGTTAATGCCATCACCTCTGACGATGTCATCAATGCTTCTGAGGCGGCGGGCACGGTGGCAGTCTCCGGCACCGCGACGGGTGGCGACATCGCAGAGGGTGATACGGTTACCCTGGAAATCAATGGTGAGACGTATACCACGACGGTGGACGCGAACGGCGAATGGTCCGTGGACGTCGCGGGCTCCGATTTGGCAGCGGATACCGCTTTCGATGCCGTCGTGACGTCCTCCGATGCGGCCGGCAACACCGTTGATACGACCGGCTCTTCCACGCATACGGTGGACACCGAAGCTACGGCAGGTACCGTTACCGTTAACGCGATCACGAGCGATGACGTGATCAATGCTTCTGAGGCAGCAGGCACGGTGGCGGTCTCTGGCACGGCGACCGGTGGTGACATTGCCGAAGGTGATACCGTTACCCTGGAGATTAACGGTGAAACCTACACCACCACCGTGGATGCAAACGGCGAGTGGAGCGTTGACGTTGCTGGCTCTGACTTGGCAGCTGACACCGCGTTTGACGCGGTCGTGACGTCTTCCGACGCAGCAGGCAACACCGTCGATACCACCGGCAGCAGCACGCACACCGTTGATACGGAAGCGACGGCGGGTACCGTGACGGTGAATGCCATCACGTCTGACGATGTTATTAACGCTTCCGAAGCGGCTGGTACGGTGGCAGTTTCCGGCACCGCGACGGGCGGCGATATCGCAGAAGGTGACACGGTTACCTTGGAAATCAACGGTGAGACGTATACCACCACGGTGGACGCCAACGGCGAGTGGTCCGTCGATGTGGCCGGCTCCGACTTGGCCGCGGATACGGCATTTGACGCCGTGGTCACCTCTTCTGATGCGGCGGGCAACACGGTTGATACCACGGGCTCCTCTACCCACACGGTTGATACCGAGGCGACCGCTGGCACCGTGACGGTGAACGCAATCACCAGCGACGATGTCATTAATGCGAGCGAAGCTGCCGGCACGGTTGCGGTTAGCGGTACCGCGACCGGTGGCGATATCGCGGAGGGTGATACGGTTACGCTGGAAATTAACGGTGAAACGTATACGACGACGGTTGACGCGAACGGTGAATGGAGCGTTGACGTTGCCGGCAGCGATTTGGCTGCGGACACGGCATTCGATGCGGTCGTTACGTCCTCTGACGCCGCTGGTAATACGGTTGACACGACGGGCTCTTCCACCCACACGGTTGACACCGAAGCGACGGCGGGTACCGTGACGGTGAATGCTATTACGAGCGATGATGTCATCAATGCTTCCGAAGCCGCTGGCACCGTGGCGGTCTCTGGCACCGCTACGGGTGGCGACATCGCGGAGGGCGACACCGTGACGCTGGAGATCAATGGCGAAACCTACACGACGACGGTGGATGCTAACGGTGAGTGGTCTGTCGATGTGGCGGGTTCCGATTTGGCGGCCGATACGGCGTTCGACGCCGTCGTGACGAGCAGCGACGCGGCCGGCAATACCGTTGATACCACGGGTAGCAGCACGCACACGGTGGATACGGAAGCCACCGCCGGTACCGTCACCGTGAATGCCATTACCTCTGATGACGTTATTAATGCGAGCGAGGCTGCGGGTACGGTTGCGGTCAGCGGCACCGCGACCGGCGGTGACATTGCAGAGGGTGATACCGTTACCTTGGAGATCAATGGTGAAACGTATACGACGACGGTGGATGCGAATGGCGAATGGTCTGTGGATGTGGCTGGCAGCGACCTGGCGGCCGACACCGCGTTCGACGCGGTGGTCACCAGCTCTGACGCTGCAGGTAATACGGTCGATACCACGGGCAGCTCTACCCATACGGTGGATACCGAAGCTACGGCCGGTACGGTTACCGTGAACGCAATCACGTCTGACGATGTTATTAACGCCTCTGAAGCAGCCGGTACCGTTGCAGTCAGCGGCACGGCTACCGGTGGCGATATTGCGGAAGGCGACACCGTTACGTTGGAGATTAACGGCGAAACGTATACGACGACCGTGGATGCCAATGGTGAATGGTCTGTGGATGTTGCCGGCTCCGATTTGGCGGCGGATACGGCCTTTGACGCTGTCGTTACCTCTAGCGACGCTGCCGGTAACACCGTGGACACGACCGGCTCCTCTACCCATACGGTCGATACGGAAGCAACGGCGGGTACGGTGACGGTTAACGCAATCACGTCCGACGACGTCATTAATGCGTCCGAGGCTGCGGGTACCGTTGCGGTGTCTGGCACCGCTACGGGCGGTGACATTGCCGAAGGTGACACGGTTACCTTGGAAATCAATGGCGAGACCTACACCACGACGGTTGATGCGAATGGCGAATGGTCTGTTGATGTTGCTGGTTCTGATCTGGCAGCCGACACCGCGTTCGATGCAGTCGTCACCAGCAGCGATGCCGCAGGCAACACGGTTGACACGACGGGCAGCAGCACGCATACCGTGGACACGGAGGCTACCGCAGGTACGGTTACGGTCAACGCCATCACCTCTGACGATGTCATCAACGCAAGCGAAGCCGCAGGCACGGTTGCAGTGTCCGGCACCGCCACCGGCGGCGATATTGCTGAAGGTGATACGGTCACGCTGGAAATCAATGGCGAGACCTACACGACCACGGTGGACGCTAATGGCGAGTGGTCTGTTGATGTGGCAGGTTCCGATTTGGCGGCAGACACCGCATTCGATGCGGTCGTTACCAGCAGCGACGCGGCAGGTAATACCGTCGATACCACGGGTTCCAGCACGCACACCGTCGATACCGAGGCTACCGCAGGTACCGTGACCGTGAACGCTATCACGTCTGATGACGTGATTAACGCGTCTGAAGCGGCCGGCACGGTGGCAGTCTCCGGTACCGCTACCGGTGGCGACATTGCGGAGGGCGATACCGTTACCCTGGAAATCAACGGTGAGACCTATACGACGACCGTGGATGCTAATGGTGAGTGGTCCGTGGACGTGGCGGGCAGCGATTTGGCCGCGGACACGGCGTTCGACGCTGTGGTCACCTCTTCCGATGCGGCCGGCAATACCGTGGACACGACGGGTTCTTCTACCCATACCGTTGACACGGAGGCGACGGCGGGCACGGTTACGGTTAACGCGATCACGTCTGATGACGTTATTAACGCAAGCGAGGCAGCTGGTACCGTCGCAGTCTCCGGCACGGCTACGGGCGGTGACATCGCTGAAGGCGATACGGTCACGTTGGAAATTAACGGCGAAACCTATACCACGACCGTTGATGCTAATGGTGAGTGGAGCGTCGATGTTGCCGGCTCCGATCTGGCGGCTGACACCGCGTTTGATGCGGTGGTGACGTCTTCTGACGCGGCCGGTAACACCGTGGATACCACGGGCTCCTCTACGCATACGGTCGACACGGAAGCTACGGCGGGTACCGTTACCGTGAACGCGATCACCTCTGATGATGTGATCAATGCATCTGAGGCTGCAGGTACCGTCGCAGTCAGCGGCACGGCGACCGGCGGCGATATCGCCGAGGGCGATACCGTGACGCTGGAAATCAATGGCGAAACGTACACCACGACGGTTGACGCAAACGGCGAGTGGAGCGTGGACGTGGCAGGCTCCGATCTGGCGGCGGATACCGCGTTTGACGCAGTCGTGACGTCTAGCGACGCCGCGGGCAATACGGTGGATACGACGGGATCCTCAACGCATACTGTTGATACGGAGGCGACGGCGGGCACGGTCACGGTGAACGCGATTACGAGCGACGACACGATTGATGGTATTGAACTGGGTCAGACGATTTCCATTTCCGGCAAAGCAGTGGGTGGTGACATCAGCGTTGGTGACGTGGTTAAGATGACGATTAATAACACCGAGTATTCCACCACCGTCAAAGCGGGTGGTATTTGGATGATCGCGGGTGTTTTGGGTTCCGATCTGGCGGCGGACTCTGAGTTTGATGTGGTGGTTACGTCCAGCGATGCTGCTGGCAACAAAGTGCAATCCATTGGCACGTCCACCCATTCCGTGGATCTGTCTGCGGAAGCGAACTTCTCTCTGGCCGAAGGCCAGCAGCATGTGTTGACCAACCTGCCGGAAGGCTTTGGTTTCCCGGACGGTACGACCGAGGTTGTTACGAATTTTGGCGGTACCATCACCCTGGGTGATGATGGCGAATATCGTTATGACGCACCGGTTCGTGACCACGGTGATGCGGTGTCCGACAAAGATTCCGTGACGGTGACCTTGGAAGATGGCCGTACGTTTACCGTGAACCTGGATATTCAAGACTCCGCACCGGTGGCGGTCGATGACCAAGATTCCATTGTCGTGCAACATGAGGAATTCGAAGTGAGCGAGATTGCGGCCAGCTGGGTGTCCTACACGCACGGCGAGTCTGTGACCACGTTTGATGGCACGTCTGATCTGGGTGGTGTGGATAACGATTCCGCGAAGGATCAGATCCGTTGGGGTAACCCGGCGGAGTCTAAGCAATCCGGCTATGGTTTCATTGACAACGATTCCAATTTGGAGGGTCGTTTTGACCTGAACCAAGACATTTCCGTGGGTACCTTCACGCACTATAACTACCCGGTGTACTCCGGCGGTGCGATTACGTCCGCAGAGATGTCCGTGGAGTTTTCCGTGCTGGACCATTTGGGTGTTAGCACGCCGGTGACCCTGACCGTGAATTTCGATCATAATGAAACCCCGAATACGAACGACGTGAACGCGTCTCGTGACATCGTCACCGTTCAAAACACCCATGTCACCTTTGAGCGCGACGGTGATATCTACACCGTGCAAATCGTGGGTTTCCGCGAGGTTGGTAACCCGGACGGCGAGGTGGTGACGTCTATCTATACGAACGAAAACGCGGCAACGTCCTACGAGCTCGTGGTCCGTGTGGTTGAAGGCGATGGTTATAGCCTGCCGTCCACGGAGGGTAACATTTTTGACGATAACGGCCTGGGTGCAGATAGCCTGGGTGCAGACGGTAGCGTGACGGTGGTTGGCGTGGCGGTTGGTGCCATTGTTTCCAGCAACGAAAGCGTGGGTCACTCCATCGAGGGCCAATATGGCAACCTGGTGCTGAACTCCGACGGTAGCTATGTTTATGACGTGACCGCATCCGTTAGCGATATCCCGGCGGGTGCCACCGAGTCTTTCGCGTATCTGATCCAGGATCAGGATGGTTCCACCAGCAGCGCAAATCTGTCCATCAACGTGGGTACGAATACCGCACCGAAAGCGCAGGACGACTCTACGCCGGATTCTTTGTTTGCAGGTCTGGTTGGCGAATACTATGGCACGAATAGCCAGTTGAACAATATTTCCGATTTTCGTGCGTTGGTGGACTCCAAAGAGGCGGACGCAACCTTTGAAGCGGCGAATATTAGCTACGGCCGTGGTTCTTCCGACGTGGCGAAGGGTACGCATTTGCAAGAGTTCTTGGGTTCTGACGCATCTACCCTGAGCACCGACCCGGGTGACAACACCGATGGCGGCATTTACCTGCAGGGTTACGTGTACTTGGAAGCGGGCACCTATAACTTCAAAGTCACCGCAGATGACGGTTACGAGATTACCATCAACGGCAATCCGGTGGCGACCGTTGACAACAACCAGTCCGTCTACACCGTGACCCACGCGTCTTTCACCATCTCTGAATCTGGTTACCAAGCAATCGACATGATCTGGTGGGATCAAGGTGGCGATTACGTGTTCCAACCGACCCTGAGCGCGGACGGTGGCTCTACCTACTTCGTTCTGGACTCTGCGATCTTGTCCTCCACGGGCGAAACGCCGTACACGACCGCGCAAGAACAAGCGTTGGAAATTAACGTGGATCGTTTGTTGGACAATGATACGGACAGCGATAACGGTGACACGTTGTCCGTGACGTCCATCTCCAATGTGAAGAACGGCTATGCATACCTGGACGCGGAAGGCATCATTCATTTTACGCCGGTCAAGGGCTTTGCAGGCGTGGCGACCATTGATTACACGATTGAGGATGGTAATGGCGGTCGTGACACCGCGACGGTGTCCATCGACGTCACGCCGGAGTCCATCCTGCCGATGGTTACGGTGAACGTTTCTCAATCCAACTCCTTCGGCTTTTGGGATGGCACGTCTACCCAGGCAGAAATCACCCATTCCTTCGATCATTACATCGGTTCCGCATTTGACGCAAGCAATAACAACGTGGCAGTGACGGGTAATGTTAGCGCGACCTTGAACGTGCTGGCGGGTGATGACAAAGTGTCTATCGATGGCAACGTGGAGGATGTTCTGGTGGCAGCGAACGTCGCGGTGCTGGATATGGGCACGGGCAATGACCAGCTGTATGTTGCGGGTGACGTGCTGGGCAAAATTGATGCAGGTACGGGCAATGATGAGATTTATATTAAAGGTGACGTGTCTGCGGCAGTCGATGCAGGCACCGGCAATGACGAAGTGTATATTGGCGGCAACCTGTCCGGTGACCTGGATGCAGGTACCGACAATGATAACATCCAAATCGGTGGTGACGTGAATGCAGCACTGAACGCGGGTACGGGCAACGATAACTTGATCATCGGCCACGATGTGTCCGGCATCGTTAACATGGGTACGGACAACGATACCGTGGAAGTGGGCCGTACCATCAATGCAAGCGGTAAAGTTCTGCTGGACACGGGCGACGACAGCCTGCTGGTGTCCGGCGACTTGTTCGGCGAAGTGGATGGCGGCACCGGTAACGATACCATCATTATCGCGGGTAAGGTTTCCGGCAACATCCAGGGTGGCACGGGCAATGACATTGTGCGTGTGCAGAGCCAGGTTTGGGCGGAGGCGAACATCTCCCTGGGTACCGGTGACGATGTGCTGATCGTGGAACACGAACTGCATGGTACCGTCGCTGGTAATGAGGGCGATGATAGCATTTACCTGAAGTTCTACACCAAAGAACAATATAACAACAACTCTGATTTGCGCAACCGCGTGGCCAATTTCGAGCACATCCGTGTGTCTGACGGCGTTGTGAAGGGTTCCCCGGCAGACTTTGCGGACTACAAGCACGGTGGCTATAGCTACGACGTTAGCGTGTCCATTGACACCAGCAATGCCAACGTTGGTGCATCCACGGTGACGTTGCTGGGTATTCCGATCGCGGGCGTTACCCTGATGCTGGCAGGTCAACCGCTGACGGCGAATGCAACCGGTGGCTACGATATCGAGGTGTCTTCCGACCAAACCGCGATTGGTGGTCTGAAGCTTGGCAATAGCCTGGCGAAAAATGTGCTGAGCGGCGGCAAGGGTAATGACAAACTGTATGGTAGCGAGGGTGCGGACCTGCTGGATGGTGGTGAGGGCAACGACCTGCTGAAGGGTGGCTACGGTAACGATATCTACCGTTATCTGAGCGGTTATGGCCACCACATCATTGACGATGAAGGTGGCAAGGACGATAAACTGAGCCTGGCGGACATCGATTTCCGTGACGTGGCGTTTAAGCGTGAGGGTAACGATCTGATTATGTACAAAGCGGAAGGCAACGTTCTGAGCATCGGTCACAAAAACGGCATTACCTTCAAGAACTGGTTTGAGAAAGAAAGCGACGATCTGAGCAACCACCAGATCGAGCAAATTTTCGACAAGGATGGTCGTGTGATTACCCCGGACAGCCTGAAGAAAGCGTTTGAATACCAGCAAAGCAACAACAAAGTGAGCTACGTTTATGGTCACGACGCGAGCACCTATGGCAGCCAGGATAACCTGAACCCGCTGATCAACGAGATTAGCAAGATCATTAGCGCGGCGGGCAACTTCGATGTTAAAGAGGAACGTAGCGCGGCGAGCCTGCTGCAACTGAGCGGTAATGCGAGCGATTTCAGCTATGGCCGTAACAGCATCACCCTGACCGCGAGCGCGCTCGAG

mRuby2-*Mp*A

ATGGTGTCGAAGGGCGAAGAGCTGATAAAAGAGAATATGCGTATGAAAGTGGTCATGGAAGGCAGTGTTAATGGACACCAATTTAAATGTACAGGAGAGGGTGAAGGAAATCCCTATATGGGCACACAGACAATGAGAATCAAGGTTATTGAAGGTGGTCCCCTGCCGTTCGCTTTCGACATCTTGGCTACCAGCTTCATGTATGGGTCCCGCACATTCATTAAGTATCCAAAGGGAATACCGGATTTTTTTAAACAGTCATTTCCTGAAGGTTTTACCTGGGAGCGAGTAACTCGTTACGAGGACGGTGGGGTCGTCACTGTGATGCAGGATACCTCCCTGGAAGATGGATGCCTGGTATATCATGTGCAAGTACGCGGCGTTAATTTTCCAAGCAATGGTCCGGTTATGCAGAAGAAAACGAAAGGTTGGGAACCTAATACGGAAATGATGTATCCAGCAGATGGTGGATTGCGCGGGTATACGCACATGGCCCTGAAAGTCGATGGAGGTGGACATTTATCATGTTCATTTGTTACTACATATAGAAGCAAAAAAACGGTCGGCAATATAAAAATGCCTGGGATCCACGCGGTGGACCACAGATTAGAGCGTTTAGAAGAATCCGACAATGAAATGTTTGTCGTTCAGCGCGAACATGCGGTTGCCAAATTTGCCGGTCTCGGTGGTGGCATGGATGAGCTCTACAAAGCTAGCAGCGACAATCAGTTTCCGTTCGCCACCTTGGGCAATGCAATCGGCTTTATCACGAAGTTGGATGGTAGCGTCACCGTTCAGAGCATCGACGGCCAAGAACGCGTGTTGAAATTGGGCGACCCGATTTTCTTTGGCGAAACCGTGTTGACCGGTGGTTCCGGTTCTGTCACGATCGCGTTCGTTGATGGTACGGATGTCGTGATTGGTGGTGATTCCATCGTTGAAATGACCGATGAGATTTACAACACCGGCGATAACGAGGATCTGGTGGCAGACAGCTCTAGCGAGATTGATGCCCTGCAAAATGCAATCCTGGCAGGCGATGACCCGACGCTGATTCAAGACGCCCCGGCTGCCGGCAATACGCTGGCCGACCAGCAGCGCGTCGATGTTTCCATTGAACGTAACGATAATTCCGCCCAAGCTGGTTTCGGCGTGGACACCCAATCCAGCCTGCCGACCTACGGCTACGATACGGACAATGGTAATGGCGGCCAAGCGACCGAGCGCGAATACTCTGCTCCGAGCCTGAGCCGCACCTTGAACCAGTCCCCGCTGCTGATCAATCTGGATATCGATCCGGTTACCGGTGACTCTGTGATTAACGCAGCAGAAGCGGGTGGTACGGTGACCTTGACCGGTGTGGTGAACGGTGACGTTTTTAGCTCCGGTGTGGTTACGCTGGTTATCAATGGTGTGACGTACTCCACGAATGTGAATCCGAACGGCACGTGGTCTGTCTCTGTTGCAGGTTCTGATCTGAGCGCGGACTCCGATCGTATTGTTGATGCAAGCGTTGTGGTCACCAACGGTGCGGGTCAGCAAGGTACGGCTGATTCCACCGAGTCCTTTATCGTCAAGACGTCTAGCCGCGCTACCATCCGCGTTAATTCCATCACGTCCGACGACGTTGTTAATGCTGAGGAGAGCAATTCCACCATCACGGTTTCTGGCCGCGTCGGCTTGGACGCCTCCGCCGGTGATACGGTCTCTATGACGATCAATGGCACCCTGTACACCACCGTTGTGTTGGCTAACAAGACCTGGTCTGTGGGCGTGAGCGGTTCTGACTTGGCACAAGATAATTCTTTTCAGGTGTCCGTGACGGGTCAAGATTCTGCGGGTAATCCGTATGCAGGCACCACGACCAGCACCCATACGGTGGATACGTCTGCAGACGCGGGCACGGTGACCGTTAATGCTATTACCTCTGACGACGTGATCAACGCTAGCGAAGCGGCTGGTACCGTCGCCGTCTCTGGCACGGCGACGGGTGGCGATATTGCCGAAGGTGACACCGTTACGTTGGAGATCAACGGTGAGACGTACACCACGACCGTGGATGCGAATGGTGAATGGAGCGTGGATGTCGCGGGTAGCGATCTGGCTGCGGACACGGCTTTCGATGCTGTTGTTACGTCTTCCGATGCAGCCGGTAACACGGTTGATACCACCGGCTCTTCCACCCATACCGTCGACACGGAGGCAACCGCTGGTACGGTTACCGTGAATGCTATCACGTCCGATGATGTCATTAACGCATCTGAAGCAGCGGGCACCGTTGCCGTTTCCGGCACGGCAACCGGCGGCGATATCGCAGAGGGTGATACGGTGACCTTGGAAATTAATGGTGAGACCTACACCACGACGGTGGATGCCAACGGCGAATGGAGCGTTGATGTTGCAGGCAGCGACCTGGCCGCTGACACGGCGTTTGACGCTGTTGTGACCTCTTCTGATGCTGCAGGTAATACGGTTGATACGACGGGTTCCAGCACCCACACCGTTGATACCGAAGCTACGGCTGGTACCGTTACGGTTAACGCGATTACGAGCGATGACGTCATCAACGCGTCCGAAGCCGCTGGTACCGTTGCGGTTTCTGGTACCGCAACGGGCGGCGATATTGCCGAAGGCGACACGGTTACCTTGGAGATTAATGGCGAGACCTATACCACGACCGTCGATGCGAACGGCGAATGGTCTGTTGATGTGGCGGGTAGCGATCTGGCAGCTGACACGGCATTTGACGCCGTGGTTACGAGCAGCGATGCGGCCGGTAATACGGTGGACACGACGGGTTCCTCCACGCACACGGTTGATACGGAGGCGACGGCGGGCACCGTTACCGTTAATGCCATCACCTCTGACGATGTCATCAATGCTTCTGAGGCGGCGGGCACGGTGGCAGTCTCCGGCACCGCGACGGGTGGCGACATCGCAGAGGGTGATACGGTTACCCTGGAAATCAATGGTGAGACGTATACCACGACGGTGGACGCGAACGGCGAATGGTCCGTGGACGTCGCGGGCTCCGATTTGGCAGCGGATACCGCTTTCGATGCCGTCGTGACGTCCTCCGATGCGGCCGGCAACACCGTTGATACGACCGGCTCTTCCACGCATACGGTGGACACCGAAGCTACGGCAGGTACCGTTACCGTTAACGCGATCACGAGCGATGACGTGATCAATGCTTCTGAGGCAGCAGGCACGGTGGCGGTCTCTGGCACGGCGACCGGTGGTGACATTGCCGAAGGTGATACCGTTACCCTGGAGATTAACGGTGAAACCTACACCACCACCGTGGATGCAAACGGCGAGTGGAGCGTTGACGTTGCTGGCTCTGACTTGGCAGCTGACACCGCGTTTGACGCGGTCGTGACGTCTTCCGACGCAGCAGGCAACACCGTCGATACCACCGGCAGCAGCACGCACACCGTTGATACGGAAGCGACGGCGGGTACCGTGACGGTGAATGCCATCACGTCTGACGATGTTATTAACGCTTCCGAAGCGGCTGGTACGGTGGCAGTTTCCGGCACCGCGACGGGCGGCGATATCGCAGAAGGTGACACGGTTACCTTGGAAATCAACGGTGAGACGTATACCACCACGGTGGACGCCAACGGCGAGTGGTCCGTCGATGTGGCCGGCTCCGACTTGGCCGCGGATACGGCATTTGACGCCGTGGTCACCTCTTCTGATGCGGCGGGCAACACGGTTGATACCACGGGCTCCTCTACCCACACGGTTGATACCGAGGCGACCGCTGGCACCGTGACGGTGAACGCAATCACCAGCGACGATGTCATTAATGCGAGCGAAGCTGCCGGCACGGTTGCGGTTAGCGGTACCGCGACCGGTGGCGATATCGCGGAGGGTGATACGGTTACGCTGGAAATTAACGGTGAAACGTATACGACGACGGTTGACGCGAACGGTGAATGGAGCGTTGACGTTGCCGGCAGCGATTTGGCTGCGGACACGGCATTCGATGCGGTCGTTACGTCCTCTGACGCCGCTGGTAATACGGTTGACACGACGGGCTCTTCCACCCACACGGTTGACACCGAAGCGACGGCGGGTACCGTGACGGTGAATGCTATTACGAGCGATGATGTCATCAATGCTTCCGAAGCCGCTGGCACCGTGGCGGTCTCTGGCACCGCTACGGGTGGCGACATCGCGGAGGGCGACACCGTGACGCTGGAGATCAATGGCGAAACCTACACGACGACGGTGGATGCTAACGGTGAGTGGTCTGTCGATGTGGCGGGTTCCGATTTGGCGGCCGATACGGCGTTCGACGCCGTCGTGACGAGCAGCGACGCGGCCGGCAATACCGTTGATACCACGGGTAGCAGCACGCACACGGTGGATACGGAAGCCACCGCCGGTACCGTCACCGTGAATGCCATTACCTCTGATGACGTTATTAATGCGAGCGAGGCTGCGGGTACGGTTGCGGTCAGCGGCACCGCGACCGGCGGTGACATTGCAGAGGGTGATACCGTTACCTTGGAGATCAATGGTGAAACGTATACGACGACGGTGGATGCGAATGGCGAATGGTCTGTGGATGTGGCTGGCAGCGACCTGGCGGCCGACACCGCGTTCGACGCGGTGGTCACCAGCTCTGACGCTGCAGGTAATACGGTCGATACCACGGGCAGCTCTACCCATACGGTGGATACCGAAGCTACGGCCGGTACGGTTACCGTGAACGCAATCACGTCTGACGATGTTATTAACGCCTCTGAAGCAGCCGGTACCGTTGCAGTCAGCGGCACGGCTACCGGTGGCGATATTGCGGAAGGCGACACCGTTACGTTGGAGATTAACGGCGAAACGTATACGACGACCGTGGATGCCAATGGTGAATGGTCTGTGGATGTTGCCGGCTCCGATTTGGCGGCGGATACGGCCTTTGACGCTGTCGTTACCTCTAGCGACGCTGCCGGTAACACCGTGGACACGACCGGCTCCTCTACCCATACGGTCGATACGGAAGCAACGGCGGGTACGGTGACGGTTAACGCAATCACGTCCGACGACGTCATTAATGCGTCCGAGGCTGCGGGTACCGTTGCGGTGTCTGGCACCGCTACGGGCGGTGACATTGCCGAAGGTGACACGGTTACCTTGGAAATCAATGGCGAGACCTACACCACGACGGTTGATGCGAATGGCGAATGGTCTGTTGATGTTGCTGGTTCTGATCTGGCAGCCGACACCGCGTTCGATGCAGTCGTCACCAGCAGCGATGCCGCAGGCAACACGGTTGACACGACGGGCAGCAGCACGCATACCGTGGACACGGAGGCTACCGCAGGTACGGTTACGGTCAACGCCATCACCTCTGACGATGTCATCAACGCAAGCGAAGCCGCAGGCACGGTTGCAGTGTCCGGCACCGCCACCGGCGGCGATATTGCTGAAGGTGATACGGTCACGCTGGAAATCAATGGCGAGACCTACACGACCACGGTGGACGCTAATGGCGAGTGGTCTGTTGATGTGGCAGGTTCCGATTTGGCGGCAGACACCGCATTCGATGCGGTCGTTACCAGCAGCGACGCGGCAGGTAATACCGTCGATACCACGGGTTCCAGCACGCACACCGTCGATACCGAGGCTACCGCAGGTACCGTGACCGTGAACGCTATCACGTCTGATGACGTGATTAACGCGTCTGAAGCGGCCGGCACGGTGGCAGTCTCCGGTACCGCTACCGGTGGCGACATTGCGGAGGGCGATACCGTTACCCTGGAAATCAACGGTGAGACCTATACGACGACCGTGGATGCTAATGGTGAGTGGTCCGTGGACGTGGCGGGCAGCGATTTGGCCGCGGACACGGCGTTCGACGCTGTGGTCACCTCTTCCGATGCGGCCGGCAATACCGTGGACACGACGGGTTCTTCTACCCATACCGTTGACACGGAGGCGACGGCGGGCACGGTTACGGTTAACGCGATCACGTCTGATGACGTTATTAACGCAAGCGAGGCAGCTGGTACCGTCGCAGTCTCCGGCACGGCTACGGGCGGTGACATCGCTGAAGGCGATACGGTCACGTTGGAAATTAACGGCGAAACCTATACCACGACCGTTGATGCTAATGGTGAGTGGAGCGTCGATGTTGCCGGCTCCGATCTGGCGGCTGACACCGCGTTTGATGCGGTGGTGACGTCTTCTGACGCGGCCGGTAACACCGTGGATACCACGGGCTCCTCTACGCATACGGTCGACACGGAAGCTACGGCGGGTACCGTTACCGTGAACGCGATCACCTCTGATGATGTGATCAATGCATCTGAGGCTGCAGGTACCGTCGCAGTCAGCGGCACGGCGACCGGCGGCGATATCGCCGAGGGCGATACCGTGACGCTGGAAATCAATGGCGAAACGTACACCACGACGGTTGACGCAAACGGCGAGTGGAGCGTGGACGTGGCAGGCTCCGATCTGGCGGCGGATACCGCGTTTGACGCAGTCGTGACGTCTAGCGACGCCGCGGGCAATACGGTGGATACGACGGGATCCTCAACGCATACTGTTGATACGGAGGCGACGGCGGGCACGGTCACGGTGAACGCGATTACGAGCGACGACACGATTGATGGTATTGAACTGGGTCAGACGATTTCCATTTCCGGCAAAGCAGTGGGTGGTGACATCAGCGTTGGTGACGTGGTTAAGATGACGATTAATAACACCGAGTATTCCACCACCGTCAAAGCGGGTGGTATTTGGATGATCGCGGGTGTTTTGGGTTCCGATCTGGCGGCGGACTCTGAGTTTGATGTGGTGGTTACGTCCAGCGATGCTGCTGGCAACAAAGTGCAATCCATTGGCACGTCCACCCATTCCGTGGATCTGTCTGCGGAAGCGAACTTCTCTCTGGCCGAAGGCCAGCAGCATGTGTTGACCAACCTGCCGGAAGGCTTTGGTTTCCCGGACGGTACGACCGAGGTTGTTACGAATTTTGGCGGTACCATCACCCTGGGTGATGATGGCGAATATCGTTATGACGCACCGGTTCGTGACCACGGTGATGCGGTGTCCGACAAAGATTCCGTGACGGTGACCTTGGAAGATGGCCGTACGTTTACCGTGAACCTGGATATTCAAGACTCCGCACCGGTGGCGGTCGATGACCAAGATTCCATTGTCGTGCAACATGAGGAATTCGAAGTGAGCGAGATTGCGGCCAGCTGGGTGTCCTACACGCACGGCGAGTCTGTGACCACGTTTGATGGCACGTCTGATCTGGGTGGTGTGGATAACGATTCCGCGAAGGATCAGATCCGTTGGGGTAACCCGGCGGAGTCTAAGCAATCCGGCTATGGTTTCATTGACAACGATTCCAATTTGGAGGGTCGTTTTGACCTGAACCAAGACATTTCCGTGGGTACCTTCACGCACTATAACTACCCGGTGTACTCCGGCGGTGCGATTACGTCCGCAGAGATGTCCGTGGAGTTTTCCGTGCTGGACCATTTGGGTGTTAGCACGCCGGTGACCCTGACCGTGAATTTCGATCATAATGAAACCCCGAATACGAACGACGTGAACGCGTCTCGTGACATCGTCACCGTTCAAAACACCCATGTCACCTTTGAGCGCGACGGTGATATCTACACCGTGCAAATCGTGGGTTTCCGCGAGGTTGGTAACCCGGACGGCGAGGTGGTGACGTCTATCTATACGAACGAAAACGCGGCAACGTCCTACGAGCTCGTGGTCCGTGTGGTTGAAGGCGATGGTTATAGCCTGCCGTCCACGGAGGGTAACATTTTTGACGATAACGGCCTGGGTGCAGATAGCCTGGGTGCAGACGGTAGCGTGACGGTGGTTGGCGTGGCGGTTGGTGCCATTGTTTCCAGCAACGAAAGCGTGGGTCACTCCATCGAGGGCCAATATGGCAACCTGGTGCTGAACTCCGACGGTAGCTATGTTTATGACGTGACCGCATCCGTTAGCGATATCCCGGCGGGTGCCACCGAGTCTTTCGCGTATCTGATCCAGGATCAGGATGGTTCCACCAGCAGCGCAAATCTGTCCATCAACGTGGGTACGAATACCGCACCGAAAGCGCAGGACGACTCTACGCCGGATTCTTTGTTTGCAGGTCTGGTTGGCGAATACTATGGCACGAATAGCCAGTTGAACAATATTTCCGATTTTCGTGCGTTGGTGGACTCCAAAGAGGCGGACGCAACCTTTGAAGCGGCGAATATTAGCTACGGCCGTGGTTCTTCCGACGTGGCGAAGGGTACGCATTTGCAAGAGTTCTTGGGTTCTGACGCATCTACCCTGAGCACCGACCCGGGTGACAACACCGATGGCGGCATTTACCTGCAGGGTTACGTGTACTTGGAAGCGGGCACCTATAACTTCAAAGTCACCGCAGATGACGGTTACGAGATTACCATCAACGGCAATCCGGTGGCGACCGTTGACAACAACCAGTCCGTCTACACCGTGACCCACGCGTCTTTCACCATCTCTGAATCTGGTTACCAAGCAATCGACATGATCTGGTGGGATCAAGGTGGCGATTACGTGTTCCAACCGACCCTGAGCGCGGACGGTGGCTCTACCTACTTCGTTCTGGACTCTGCGATCTTGTCCTCCACGGGCGAAACGCCGTACACGACCGCGCAAGAACAAGCGTTGGAAATTAACGTGGATCGTTTGTTGGACAATGATACGGACAGCGATAACGGTGACACGTTGTCCGTGACGTCCATCTCCAATGTGAAGAACGGCTATGCATACCTGGACGCGGAAGGCATCATTCATTTTACGCCGGTCAAGGGCTTTGCAGGCGTGGCGACCATTGATTACACGATTGAGGATGGTAATGGCGGTCGTGACACCGCGACGGTGTCCATCGACGTCACGCCGGAGTCCATCCTGCCGATGGTTACGGTGAACGTTTCTCAATCCAACTCCTTCGGCTTTTGGGATGGCACGTCTACCCAGGCAGAAATCACCCATTCCTTCGATCATTACATCGGTTCCGCATTTGACGCAAGCAATAACAACGTGGCAGTGACGGGTAATGTTAGCGCGACCTTGAACGTGCTGGCGGGTGATGACAAAGTGTCTATCGATGGCAACGTGGAGGATGTTCTGGTGGCAGCGAACGTCGCGGTGCTGGATATGGGCACGGGCAATGACCAGCTGTATGTTGCGGGTGACGTGCTGGGCAAAATTGATGCAGGTACGGGCAATGATGAGATTTATATTAAAGGTGACGTGTCTGCGGCAGTCGATGCAGGCACCGGCAATGACGAAGTGTATATTGGCGGCAACCTGTCCGGTGACCTGGATGCAGGTACCGACAATGATAACATCCAAATCGGTGGTGACGTGAATGCAGCACTGAACGCGGGTACGGGCAACGATAACTTGATCATCGGCCACGATGTGTCCGGCATCGTTAACATGGGTACGGACAACGATACCGTGGAAGTGGGCCGTACCATCAATGCAAGCGGTAAAGTTCTGCTGGACACGGGCGACGACAGCCTGCTGGTGTCCGGCGACTTGTTCGGCGAAGTGGATGGCGGCACCGGTAACGATACCATCATTATCGCGGGTAAGGTTTCCGGCAACATCCAGGGTGGCACGGGCAATGACATTGTGCGTGTGCAGAGCCAGGTTTGGGCGGAGGCGAACATCTCCCTGGGTACCGGTGACGATGTGCTGATCGTGGAACACGAACTGCATGGTACCGTCGCTGGTAATGAGGGCGATGATAGCATTTACCTGAAGTTCTACACCAAAGAACAATATAACAACAACTCTGATTTGCGCAACCGCGTGGCCAATTTCGAGCACATCCGTGTGTCTGACGGCGTTGTGAAGGGTTCCCCGGCAGACTTTGCGGACTACAAGCACGGTGGCTATAGCTACGACGTTAGCGTGTCCATTGACACCAGCAATGCCAACGTTGGTGCATCCACGGTGACGTTGCTGGGTATTCCGATCGCGGGCGTTACCCTGATGCTGGCAGGTCAACCGCTGACGGCGAATGCAACCGGTGGCTACGATATCGAGGTGTCTTCCGACCAAACCGCGATTGGTGGTCTGAAGCTTGGCAATAGCCTGGCGAAAAATGTGCTGAGCGGCGGCAAGGGTAATGACAAACTGTATGGTAGCGAGGGTGCGGACCTGCTGGATGGTGGTGAGGGCAACGACCTGCTGAAGGGTGGCTACGGTAACGATATCTACCGTTATCTGAGCGGTTATGGCCACCACATCATTGACGATGAAGGTGGCAAGGACGATAAACTGAGCCTGGCGGACATCGATTTCCGTGACGTGGCGTTTAAGCGTGAGGGTAACGATCTGATTATGTACAAAGCGGAAGGCAACGTTCTGAGCATCGGTCACAAAAACGGCATTACCTTCAAGAACTGGTTTGAGAAAGAAAGCGACGATCTGAGCAACCACCAGATCGAGCAAATTTTCGACAAGGATGGTCGTGTGATTACCCCGGACAGCCTGAAGAAAGCGTTTGAATACCAGCAAAGCAACAACAAAGTGAGCTACGTTTATGGTCACGACGCGAGCACCTATGGCAGCCAGGATAACCTGAACCCGCTGATCAACGAGATTAGCAAGATCATTAGCGCGGCGGGCAACTTCGATGTTAAAGAGGAACGTAGCGCGGCGAGCCTGCTGCAACTGAGCGGTAATGCGAGCGATTTCAGCTATGGCCGTAACAGCATCACCCTGACCGCGAGCGCGCTCGAG

HlyAc

GGCAATAGCCTGGCGAAAAATGTGCTGAGCGGCGGCAAGGGTAATGACAAACTGTATGGTAGCGAGGGTGCGGAC

CTGCTGGATGGTGGTGAGGGCAACGACCTGCTGAAGGGTGGCTACGGTAACGATATCTACCGTTATCTGAGCGGT

TATGGCCACCACATCATTGACGATGAAGGTGGCAAGGACGATAAACTGAGCCTGGCGGACATCGATTTCCGTGAC

GTGGCGTTTAAGCGTGAGGGTAACGATCTGATTATGTACAAAGCGGAAGGCAACGTTCTGAGCATCGGTCACAAA

AACGGCATTACCTTCAAGAACTGGTTTGAGAAAGAAAGCGACGATCTGAGCAACCACCAGATCGAGCAAATTTTC

GACAAGGATGGTCGTGTGATTACCCCGGACAGCCTGAAGAAAGCGTTTGAATACCAGCAAAGCAACAACAAAGTG

AGCTACGTTTATGGTCACGACGCGAGCACCTATGGCAGCCAGGATAACCTGAACCCGCTGATCAACGAGATTAGC

AAGATCATTAGCGCGGCGGGCAACTTCGATGTTAAAGAGGAACGTAGCGCGGCGAGCCTGCTGCAACTGAGCGGT

AATGCGAGCGATTTCAGCTATGGCCGTAACAGCATCACCCTGACCGCGAGCGCG

Lysostaphin-HlyAc (glycine-rich cleavable linker)

ATGACCTCAAAGGCCCTTGTGCAAAATCGTACAGCGTTACGCGCAGCGACACACGAACATTCTGCTCAGTGGCTGAATAATTATAAAAAAGGTTATGGTTACGGTCCCTACCCGCTGGGAATCAATGGAGGAATGCACTACGGTGTGGACTTCTTTATGAACATTGGCACACCGGTCAAGGCGATTAGTTCAGGTAAAATCGTAGAAGCTGGTTGGTCAAACTATGGCGGTGGGAATCAAATCGGTCTTATCGAGAATGATGGGGTGCATCGTCAGTGGTACATGCACCTGAGCAAATATAATGTGAAAGTTGGCGACTATGTAAAGGCCGGGCAAATCATTGGGTGGAGCGGCTCTACTGGTTATTCTACAGCTCCGCATCTTCATTTCCAACGCATGGTTAATAGTTTTTCAAACTCTACAGCCCAAGACCCTATGCCCTTTTTGAAGTCTGCGGGTTACGGGAAGGCCGGTGGAACCGTTACCCCCACGCCAAATACCGGCTGGAAAACGAATAAGTATGGAACACTGTATAAGTCTGAATCCGCCTCGTTCACTCCCAACACGGATATTATCACACGCACCACTGGACCATTCCGTTCCATGCCTCAGTCAGGCGTTTTAAAGGCGGGTCAGACAATCCACTACGACGAGGTTATGAAGCAGGACGGACATGTCTGGGTCGGGTACACGGGTAACTCCGGGCAACGTATCTACTTGCCGGTCCGCACGTGGAATAAAAGCACCAATACACTGGGCGTACTTTGGGGCACAATTAAGGGAGGGGGCGGATCGCTGGTTCCTCGCGGGTCAGGAGGGGGCGGGAGCGGAAATTCTCTTGCTAAAAATGTATTATCCGGTGGAAAAGGTAATGACAAGTTGTACGGCAGTGAGGGAGCAGACCTGCTTGATGGCGGAGAAGGGAATGATCTTCTGAAAGGTGGATATGGTAATGATATTTATCGTTATCTTTCAGGATATGGCCATCATATTATTGACGATGAAGGGGGGAAAGACGATAAACTCAGTTTAGCTGATATAGATTTCCGGGACGTTGCCTTTAAGCGAGAAGGGAATGACCTCATTATGTATAAAGCTGAAGGTAATGTTCTTTCTATTGGCCACAAAAATGGTATTACATTTAAAAACTGGTTTGAAAAAGAGTCAGATGATCTCTCTAATCATCAGATAGAGCAGATTTTTGATAAAGACGGCAGGGTAATCACACCAGATTCTCTTAAAAAAGCATTTGAATATCAGCAGAGTAATAACAAGGTAAGTTATGTGTATGGACATGATGCATCAACTTATGGGAGCCAGGACAATCTTAATCCATTAATTAATGAAATCAGCAAAATCATTTCAGCTGCAGGTAACTTCGATGTTAAGGAGGAAAGATCTGCCGCTTCTTTATTGCAGTTGTCCGGTAATGCCAGTGATTTTTCATATGGACGGAACTCAATAACTTTGACAGCATCAGCACACCACCACCACCACCACTGA

HlyB/D DNA sequence:

**GATTCTTGTCATAAAATTGATTATGGGTTATACGCCCTGGAGATTTTAGCCCAATACCATAACGTCTCTGTTAACCCGGAAGAAATTAAACATAGATTTGACACAGACGGGACTGGTCTGGGATTAACGTCATGGTTGCTTGCTGCGAAATCTTTAGAACTAAAGGTAAAACAGGTAAAAAAAACAATTGACCGATTAAACTTTATTTCTCTGCCCGCATTAGTCTGGAGAGAGGATGGACGTCATTTTATTCTGACTAAAGTCAGTAAAGAAGCAAACAGATATCTTATTTTTGATCTGGAGCAGCGAAATCCCCGTGTTCTCGAACAGTCTGAGTTTGAGGCGTTATATCAGGGGCATATTATTCTTATCGCTTCCCGTTCTTCTGTTGCCGGGAAACTGGCGAAATTTGACTTTACCTGGTTTATTCCTGCCATTATAAAATACAGGAGAATATTTATTGAAACCCTTGTTGTGTCTGTTTTTTTACAATTATTTGCATTAATAACCCCCCTTTTTTTTCAGGTGGTTATGGACAAAGTATTAGTGCACAGGGGATTTTCAACTCTTAATGTTATTACTGTCGCATTATCTGTTGTGGTGGTGTTTGAGATTATACTCAGCGGTTTAAGAACTTACATTTTTGCACATAGTACAAGTCGGATTGATGTTGAGTTGGGTGCCAAACTCTTCCGGCATTTACTGGCGCTACCGATCTCTTATTTTGAGAGTCGTCGTGTTGGTGATACTGTTGCCAGGGTAAGAGAATTAGACCAGATCCGTAATTTTCTGACAGGACAGGCATTAACATCTGTTCTGGACTTATTATTTTCATTCATATTTTTTGCGGTAATGTGGTATTACAGTCCAAAGCTTACTCTGGTGATCTTATTTTCGCTGCCTTGTTATGCTGCATGGTCTGTTTTTATTAGCCCCATTTTGCGACGTCGCCTTGATGATAAGTTTTCACGGAATGCGGATAATCAATCTTTCCTGGTGGAATCAGTCACGGCGATTAACACTATAAAAGCTATGGCAGTCTCACCTCAGATGACGAACATATGGGACAAACAATTGGCAGGATATGTTGCTGCAGGCTTCAAAGTGACAGTATTAGCAACCATTGGTCAACAAGGAATACAGTTAATACAAAAGACTGTTATGATCATCAACCTGTGGTTGGGAGCACACCTGGTTATTTCCGGGGATTTATCGATTGGTCAGTTAATTGCTTTTAATATGCTTGCTGGTCAGATTGTTGCACCGGTTATTCGCCTTGCACAAATCTGGCAGGATTTCCAGCAGGTTGGTATATCAGTTACCCGCCTTGGTGATGTGCTTAACTCTCCAACTGAAAGTTATCATGGGAAACTGGCATTACCGGAAATTAATGGTGATATCACTTTTCGTAATATCCGGTTTCGCTATAAGCCTGACTCTCCGGTTATTTTAGATAATATCAATCTCAGTATTAAGCAGGGGGAGGTTATTGGTATTGTCGGACGTTCTGGTTCAGGAAAAAGCACATTAACTAAATTAATTCAACGTTTTTATATTCCTGAAAATGGCCAGGTCTTAATTGATGGACATGATCTTGCGTTGGCCGATCCTAACTGGTTACGTCGTCAGGTGGGGGTTGTGTTGCAGGACAATGTGCTGCTTAATCGCAGTATTATTGATAATATCTCACTGGCTAATCCTGGTATGTCCGTCGAAAAAGTTATTTATGCAGCGAAATTAGCAGGCGCTCATGATTTTATTTCTGAATTGCGTGAGGGGTATAACACCATTGTCGGGGAACAGGGGGCAGGATTATCCGGAGGTCAACGTCAACGCATCGCAATTGCAAGGGCGCTGGTGAACAACCCTAAAATACTTATTTTTGATGAAGCAACCAGTGCTCTGGATTATGAGTCGGAGCATATCATCATGCGCAATATGCACAAAATATGTAAGGGCAGAACGGTTATAATCATTGCTCATCGTCTGTCTACAGTAAAAAATGCAGACCGCATTATTGTCATGGAAAAAGGGAAAATTGTTGAACAGGGTAAACATAAGGAACTGCTTTCTGAACCGGAAAGTTTATACAGTTACTTATATCAGTTACAGTCAGACTAACAGAAAGAACAGAAGAATATGAAAACATGGTTAATGGGGTTCAGCGAGTTCCTGTTGCGCTATAAACTTGTCTGGAGTGAAACATGGAAAATCCGGAAGCAATTAGATACTCCGGTACGTGAAAAGGACGAAAATGAATTCTTACCCGCTCATCTGGAATTAATTGAAACGCCGGTATCCAGGCGGCCGCGTCTGGTTGCTTATTTTATTATGGGGTTTCTGGTTATTGCTGTCATTTTATCTGTTTTAGGTCAGGTGGAAATTGTTGCCACTGCAAATGGGAAATTAACACTAAGTGGGCGTAGCAAAGAAATTAAACCTATTGAAAACTCAATAGTTAAAGAAATTATCGTAAAAGAAGGAGAGTCAGTCCGGAAAGGGGATGTGTTATTAAAGCTTACAGCACTGGGAGCTGAAGCTGATACGTTAAAAACACAGTCATCACTGTTACAGACCAGGCTGGAACAAACTCGGTATCAAATTCTGAGCAGGTCAATTGAATTAAATAAACTACCTGAACTGAAGCTTCCTGATGAGCCTTATTTTCAGAATGTATCTGAAGAGGAAGTACTGCGTTTAACTTCTTTGATAAAAGAACAGTTTTCCACATGGCAAAATCAGAAGTATCAAAAAGAACTGAATCTGGATAAGAAAAGAGCAGAGCGATTAACAATACTTGCCCGTATAAACCGTTATGAAAATTTATCCCGGGTTGAAAAAAGCCGTCTGGATGATTTCAGGAGTTTATTGCATAAACAGGCAATTGCAAAACATGCTGTACTTGAGCAGGAGAATAAATATGTCGAGGCAGCAAATGAATTACGGGTTTATAAATCGCAACTGGAGCAAATTGAGAGTGAGATATTGTCTGCAAAAGAAGAATATCAGCTTGTCACGCAGCTTTTTAAAAATGAAATTTTAGACAAGCTAAGACAAACAACAGACAGCATTGAGTTATTAACTCTGGAGTTAGAGAAAAATGAAGAGCGTCAACAGGCTTCAGTAATCAGGGCCCCTGTTTCGGGAAAAGTTCAGCAACTGAAGGTTCATACTGAAGGTGGGGTTGTTACAACAGCGGAAACACTGATGGTCATCGTTCCGGAAGATGACACGCTGGAGGTTACTGCTCTGGTACAAAATAAAGATATTGGTTTTATTAACGTCGGGCAGAATGCCATCATTAAAGTGGAGGCCTTTCCTTACACCCGATATGGTTATCTGGTGGGTAAGGTGAAAAATATAAATTTAGATGCAATAGAAGACCAGAAACTGGGACTCGTTTTTAATGTCATTGTTTCTGTTGAAGAGAATGATTTGTCAACCGGGAATAAGCACATTCCATTAAGCTCGGGTATGGCTGTCACTGCAGAAATAAAGACTGGAATGCGAAGCGTAATCAGCTATCTTCTTAGTCCTCTGGAAGAGTCTGTAACAGAAAGTTTACATGAGCGT**TAA
